# Ankle2, A Target of Zika Virus, Controls Asymmetric Cell Division of Neuroblasts and Uncovers a Novel Microcephaly Pathway

**DOI:** 10.1101/611384

**Authors:** N. Link, H. Chung, A. Jolly, M. Withers, B. Tepe, B.R. Arenkiel, P.S. Shah, N.J. Krogan, H. Aydin, B.B. Geckinli, T. Tos, S. Isikay, B. Tuysuz, G.H. Mochida, A.X. Thomas, R.D. Clark, G.M. Mirzaa, J.R. Lupski, H.J. Bellen

## Abstract

Neuroblasts in flies divide asymmetrically by establishing polarity, distributing cell fate determinants asymmetrically, and positioning their spindle for cell division. The apical complex contains aPKC, Bazooka (Par3), and Par6, and its activity depends on L(2)gl. We show that Ankle2 interacts with L(2)gl and affects aPKC. Reducing Ankle2 levels disrupts ER and nuclear envelope morphology, releasing the kinase Ballchen/VRK1 into the cytosol. These defects are associated with reduced phosphorylation of aPKC, disruption of Par complex localization, and spindle alignment defects. Importantly, removal of one copy of *ballchen/VRK1* or *l(2)gl* suppresses the loss of *Ankle2* and restores viability and brain size. The Zika virus NS4A protein interacts with *Drosophila* Ankle2 and VRK1 in dividing neuroblasts. Human mutational studies implicate this neural cell division pathway in microcephaly and motor neuron disease. In summary, NS4A, ANKLE2, VRK1 and LLGL1 define a novel pathway that impinges on asymmetric determinants of neural stem cell division.

## INTRODUCTION

Proper development of the human brain requires an exquisitely coordinated series of steps and is disrupted in disorders associated with congenital microcephaly. Congenital microcephaly is characterized by reduced brain size (using occipital frontal circumference, OFC, as a surrogate measure) more than two standard deviations below the mean (Z-score < −2) at birth. It is associated with neurodevelopmental disorders, such as developmental delay and intellectual disability (DD/ID) and can be caused by external exposures to toxins, *in utero* infections, or gene mutations. Pathogenic gene variants for microcephaly have been identified through targeted genetic testing, genomic copy number studies, and exome sequencing (ES) (Brunetti-Pierri *et al.*, 2008; Dumas *et al.*, 2012; Lupski, 2015; Shinawi *et al.*, 2010; Shaheen *et al.*, 2018), elucidating about 18 primary microcephaly loci. Many syndromes significantly overlap with classic microcephaly phenotypes, and together, these disorders can be caused by defects in a wide variety of biological processes, including centriole biogenesis, DNA replication, DNA repair, cell cycle and cytokinesis, genome stability, as well as multiple cell signaling pathways (Jayaraman, Bae and Walsh, 2018).

A forward, mosaic screen for neurodevelopmental and neurodegenerative phenotypes associated with lethal mutations on the X-chromosome in *Drosophila* identified 165 loci, many with corresponding human genetic disease trait phenotypes (Yamamoto *et al.*, 2014). Among them, a mutation in *Ankle2 (Ankryin repeat and LEM domain containing 2)* causes loss of Peripheral Nervous System (PNS) organs in adult mutant clones and severely reduced brain size in hemizygous third instar larvae. To identify patients with pathogenic variants in *ANKLE2*, we surveyed the exome database of the Baylor-Hopkins Center for Mendelian Genomics (BHCMG) (Bamshad *et al.*, 2012; Posey *et al.*, 2019) and identified compound heterozygous mutations in *ANKLE2* in two siblings. Both infants exhibited severe microcephaly (Z-score = −9), and the surviving patient displayed cognitive and neurological deficits alongside extensive intellectual and developmental disabilities. Previously, we showed that mutations in *Ankle2* lead to cell loss of neuroblasts and affected neuroblast division in the developing third instar larval brain. Remarkably, expression of the wild type human ANKLE2 in flies rescued the observed mutant phenotypes (Yamamoto *et al.*, 2014). Here we explore the molecular pathways and proteins that are affected by *Ankle2* loss.

ANKLE2 belongs to a family of proteins containing LEM (<u>L</u>AP2, <u>E</u>merin, <u>M</u>AN1) domains that typically associate with the inner nuclear membrane (Lin *et al.*, 2000; Barton, Soshnev and Geyer, 2015). Conventional LEM proteins have been shown to interact with BAF (Barrier to Autointegration Factor), which binds to both DNA and the nuclear lamina (Segura-Totten *et al.*, 2002) to organize nuclear and chromatin structure. However, the LEM domain in *Drosophila* and *C. elegans* Ankle2 is not obviously conserved (Marchler-Bauer *et al.*, 2017). Studies in *C. elegans* indicate that a homolog of ANKLE2 regulates nuclear envelope morphology and functions in mitosis to promote reassembly of the nuclear envelope upon mitotic exit (Asencio *et al.*, 2012; Snyers *et al.*, 2018). During this process, ANKLE2 modulates the activities of VRK1 (Vaccina Related Kinase 1) and PP2A (Protein Phosphatase 2A) (Asencio *et al.*, 2012). However, all experiments in worms were performed at the embryonic two-cell stage, and no other phenotypes were reported except early lethality. Whilst mutations in *ANKLE2* have been associated with severe microcephaly (OFC z-sore = −2.5 to −9), human *VRK1* pathogenic variant alleles can cause a neurological disease trait consisting of complex motor and sensory axonal neuropathy and microcephaly (Gonzaga-Jauregui *et al.*, 2013).

Mutations in both *Ankle2* and the fly homologue of *VRK1, ballchen*, cause a loss of neuroblasts in 3^rd^ instar larval brains in *Drosophila* (Yamamoto *et al.*, 2014; Yakulov *et al.*, 2014). Neuroblasts (NBs) divide asymmetrically and are often used as a model to investigate stem cell biology (Homem and Knoblich, 2012) and asymmetric cell division (Gallaud *et al*., 2017). Most NBs in the larval central brain give rise to another NB and a smaller ganglion mother cell (GMC), which then divides once again to produce neurons or glia. Proper NB maintenance and regulation is essential for precise development of the adult nervous system, and misregulation of NB number or function can lead to defects in brain size (Wang *et al.*, 2009; Gateff and Schneiderman, 1974).

Congenital Zika infection in humans during pregnancy has been associated with severe microcephaly that can be as dramatic as certain genetic forms of microcephaly including phenotypes associated with bi-allelic mutations in MCPH16/*ANKLE2* (Moore *et al.*, 2017; Yamamoto *et al.*, 2014). Recently, we showed that a Zika protein, NS4A, physically interacts with ANKLE2 in human cells. Expression of NS4A in larval brains causes microcephaly, induces apoptosis, and reduces proliferation. Importantly, expression of human ANKLE2 in flies that express NS4A suppresses the associated phenotypes, demonstrating that NS4A interacts with the ANKLE2 protein and inhibits its function (Shah *et al.*, 2018). Interestingly, the Zika virus crosses the blood brain barrier and targets radial glial cells, the neural progenitors in the vertebrate cortex (Devhare *et al.*, 2017; Tang *et al.*, 2016).

Here, we show that Ankle2 is localized to the endoplasmic reticulum and nuclear envelope, like NS4A (Shah *et al.*, 2018), and genetically interacts with *ball/VRK1* to regulate brain size in flies. An allelic series at the *ANKLE2* and *VRK1* loci shows that perturbation of this pathway results in neurological disease including microcephaly. Our data indicate that the Ankle2-Ball/VRK1 pathway is required for proper localization of asymmetric proteins and spindle alignment during NB cell division by affecting two proteins, aPKC and L(2)gl, that play critical roles in the asymmetric segregation of cell fate determinants. In addition, NS4A expression in neuroblasts mimics phenotypes seen in *Ankle2* mutants, and NS4A induced microcephaly is suppressed by removing a single copy of *ball/VRK1.* Human genomics variant data and disease trait correlations extend this asymmetric cell division pathway from proteins identified in flies and reveal insights into neurological disease. In summary, NS4A hijacks the Ankle2-Ball/VRK1 pathway, which regulates progenitor stem cell asymmetric division during brain development and defines a novel human microcephaly pathway.

## RESULTS

### Human ANKLE2 variants cause microcephaly

We previously reported that compound heterozygous variants in *ANKLE2* are associated with microcephaly (Z-score = −9) (MCPH16, MIM#616681) in two affected siblings (Yamamoto *et al.*, 2014). Here, we report two additional probands carrying unique variants in *ANKLE2* identified in Seattle (LR17-511 and LR18-033; **Figure 1** and **S1, Table S1**). Brain MRIs of an age matched control (**Figure 1A**) and a proband with microcephaly from family LR17-511 document one of the more severe cases of microcephaly (Z-score = −8) (**Figure 1B)**. To investigate potential genotype-phenotype correlations, we explored the spectrum of reported neurological disease trait manifestations associated with *ANKLE2* present in the Baylor Genetics (BG) Laboratories databases. These contain clinical exome sequencing (ES) of patients with presumed genetic disorders. We screened for rare biallelic variants, predicted damaging, in *ANKLE2* that fulfill Mendelian expectations for a recessive disease trait. Three families were found to fulfill these criteria in probands with neurologically associated phenotypes (**Figure 1D** and **S1, Table S2**). These cases suggest that a diverse set of variants in *ANKLE2* may be associated with a spectrum of neurologic disease (**Figure 1**) and reveal either sporadic disease, apparent vertical transmission, and in some cases, consanguineous parentage (Yamamoto *et al.*, 2014; Shaheen *et al.*, 2018); (**Figure S1**). The identified mutations are missense, nonsense, or splicing variants that lead to premature stop codons; all subjects have biallelic variants, either compound heterozygous or homozygous alleles (**Figure 1** and **S1**). Probands exhibit congenital microcephaly (**Figure 1D**), but some also present with severe brain MRI abnormalities and skin pigmentation abnormalities (**Figure 1D**). These aggregate data demonstrate that mutations in *ANKLE2* cause autosomal recessive microcephaly.

**Figure 1:**
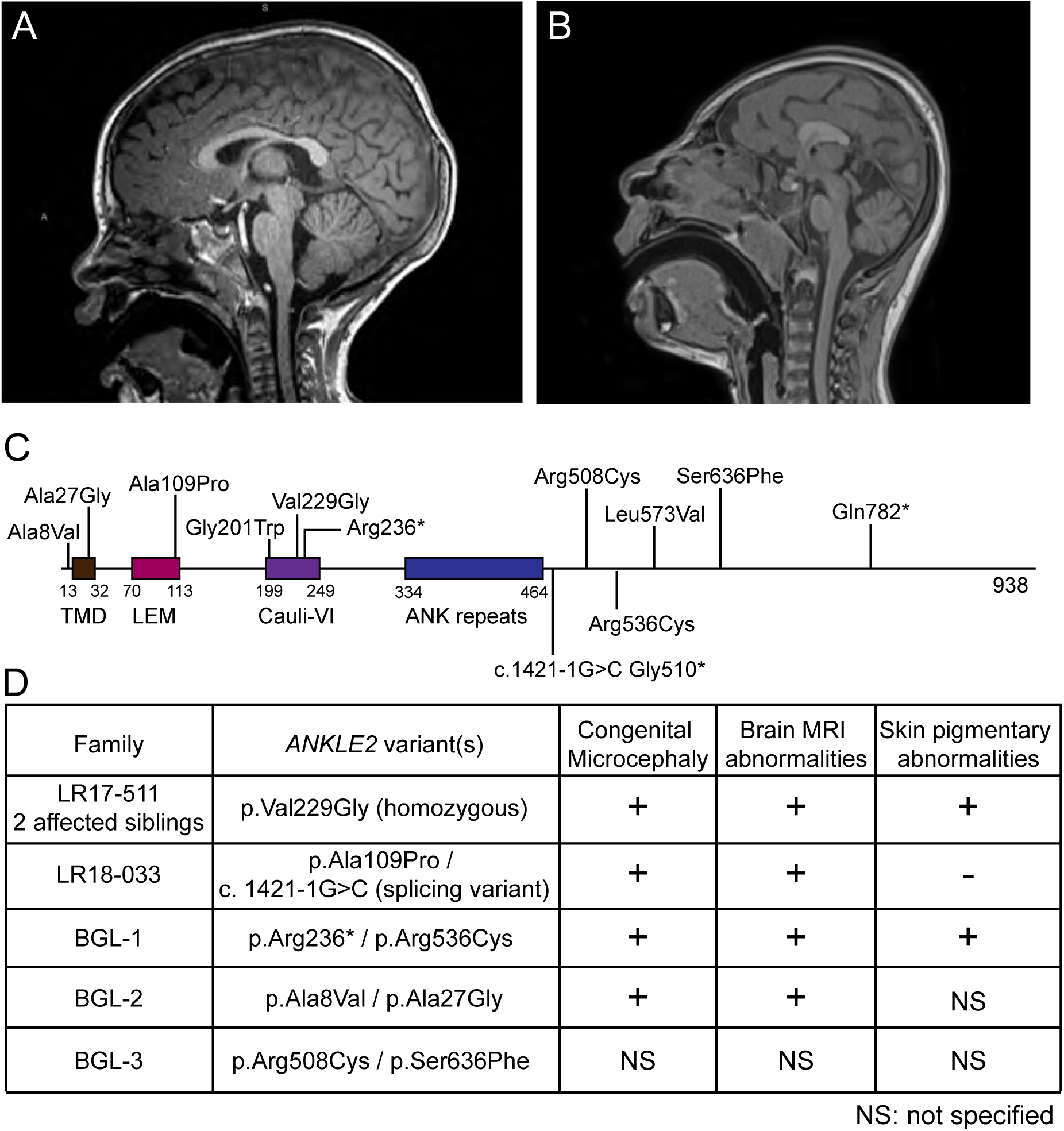
Human ANKLE2 variants in microcephaly patients. (A) Normal T1 brain MRI of a two year old infant with no microcephaly. (B) A T1 midsagittal image documents a severe microcephaly with a low-sloping forehead and partial agenesis of the corpus callosum. (C) ANKLE2 protein with known domains and the known patient variants described in this manuscript (**Figure 2**). TMD=transmembrane domain. LEM=<u>L</u>ap2, <u>E</u>merin, <u>M</u>AN1 domain. Cauli-VI= Caulimovirus viroplasmin VI domain. ANK=Ankyrin repeats. (D) Probands affected with microcephaly and associated *ANKLE2* variants and phenotypes. NS=not specified. See also **Figure S1** and **Table S1-S2**

### Null alleles of *Ankle2* are associated with reduced brain size in flies

Given the human genetic implications noted above, we used *Drosophila* to elucidate molecular mechanisms underlying *ANKLE2* associated microcephaly. The mutation originally identified in flies, *Ankle2*^*A*^ (L326A), causes reduced brain size in third instar larvae and leads to pupal lethality at temperatures ≥ 22°C (**Figure S2**). It results in decreased neuroblast number, reduced cell divisions, and a high incidence of apoptotic cell death (Yamamoto *et al.*, 2014). To create a severe loss of function allele for *Ankle2*, we integrated a CRIMIC construct containing *attP-FRT-SA-3XSTOP-polyA-3xP3-EGFP-FRT-attP* sequences using CRISPR-Cas9 in the fifth intron shared by all isoforms (**Figure 2A**; pM14; (Lee *et al.*, 2018)). The polyA sequences arrest transcription, leading to a truncated transcript that likely corresponds to a null allele (*Ankle2*^*CRIMIC*^, **Figure 2A**). These animals die as 3^rd^ instar larvae (**Figure 2B**), are smaller than wild type and *Ankle2*^*A*^ animals, and show severely reduced brain volume (**Figure 2E versus I, and M**) with complete disruption of the optic lobe (**Figure 2I**).

**Figure 2:**
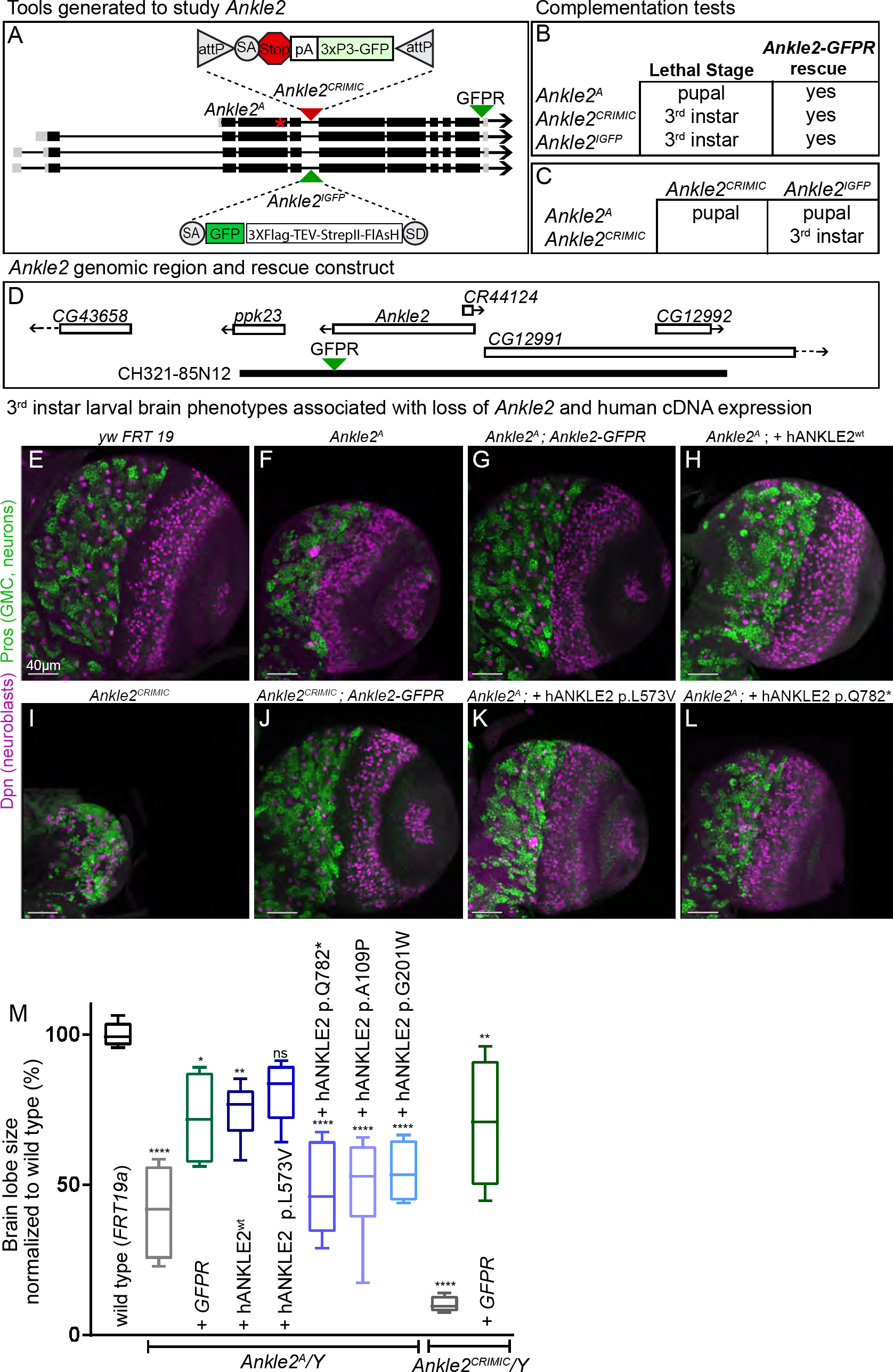
Mutations in *ANKLE2* cause microcephaly in humans and *Drosophila*. The structure and transcripts of the *Ankle2* gene with insertional mutations or tags are shown in (A). *Ankle2*^*A*^, noted by *, is an EMS generated L326H mutation, while *Ankle2*^*CRIMIC*^ represents a Crispr-Cas9 mediated MIMiC-like (CRIMIC) insertion in the 4^th^ coding intron. ΦC31 mediated cassette exchange replaced the stop-polyA in *Ankle2*^*CRIMIC*^ with an artificial exon cassette consisting of SA-GFP-SD (Venken *et al.*, 2011) to produce *Ankle2*^*IGFP*^. A C-terminal GFP tagged *Ankle2* genomic rescue construct (Ankle2-GFPR) was generated using recombineering. (B) Lethal stages and rescue by Ankle2-GFPR or (C) complementation tests of *Ankle2*^*A*^, *Ankle2*^*CRIMIC*^, *and Ankle2*^*IGFP*^. (D) *Ankle2* gene structure and genomic rescue construct (CH321-85N12). (E-L) Partial projections of 3^rd^ instar larval brains stained with Deadpan (Dpn) (purple, neuroblasts) a neuroblast marker (Bier *et al.*, 1992) and Pros (green, GMC, neurons), a neuronal lineage marker (Campbell *et al.*, 1994), to document overall brain structure is shown in (E) wild type (*y w FRT19a*), (F) *y w Ankle2*^*A*^ *FRT19a* (G) *y w Ankle2*^*A*^ *FRT19a; Ankle2-GFPR*, (H) *y w Ankle2*^*A*^ *FRT19a; da::hANKLE2*^*wt*^, (I) *Ankle2*^*CRIMIC*^, (J) *Ankle2*^*CRIMIC*^; *Ankle2-GFPR* (K) *y w Ankle2*^*A*^ *FRT19a; da:: hANKLE2 p.L573V* and (L) *y w Ankle2*^*A*^ *FRT19a; da:: hANKLE2 p.Q782\** animals. The C-terminal GFP-tagged rescue P[acman] clone CH321-85N12, (*Ankle2-GFPR*, diagramed in A and B) rescues brain morphology, brain size, and lethality in both *y w Ankle2*^*A*^ *FRT19a* (G) and *Ankle2*^*CRIMIC*^ (J) animals. Quantification of brain size is shown in (M). Box plots hinges represent the 25^th^ to 75^th^ percentiles, the central line is the median, and whiskers represent min to max. Note that hANKLE2^wt^ and hANKLE2 p.L573V rescue brain size and lethality of *Ankle2*^*A*^ mutants, whereas hANKLE2 p.Q782*, hANKLE2 p.A109P, and hANKLE2 p.G201W do not. Here, Ankle2-GFPR is reported as GFPR. See also **Figure S2**.

To determine whether Ankle2 is expressed in the brain, we used the CRIMIC allele to introduce an artificial exon that contains SA-GFP-SD in frame which produces a tagged fusion protein (*Ankle2*^*IGFP*^, **Figure 2A-B**) (Lee *et al.*, 2018). We readily detect Ankle2^IGFP^ protein in brains of heterozygous animals (**Figure S3**). However, homozygous animals are lethal and exhibit very small brains, indicating that integration of this exon disrupts protein function. Based on complementation tests, the strength of the allelic series is *Ankle2*^*A*^ < *Ankle2*^*CRIMIC*^= *Ankle2*^*IGFP*^ (**Figure 2B and C**). We therefore used recombineering (Venken *et al.*, 2006) to add a C-terminal GFP tag to Ankle2 in a BAC (CH321-85N12; referred to as *Ankle2-GFPR*, **Figure 2A and D**) (Venken *et al*., 2009). When this P[acman] clone was introduced in all three *Ankle2* mutant backgrounds, *Ankle2-GFPR* rescued brain phenotypes and lethality of these alleles (**Figure 2B, G, and J**). Hence, the chromosomes carrying the three *Ankle2* alleles do not carry second site mutations that affect brain size or viability and the tagged protein is likely to reflect the endogenous Ankle2 protein distribution.

The human reference *ANKLE2* gene rescues lethality and small brain phenotypes of *Ankle2*^*A*^ animals when expressed ubiquitously (*da-GAL4 >UAS-hANKLE2*, **Figure 2H, M**). To determine whether the microcephaly associated mutations in human ANKLE2 are loss of function alleles, we next expressed *ANKLE2 p.L573V, ANKLE2 p.Q782*, ANKLE2 p.A109P, ANKLE2 p.G201W*, in *Ankle2*^*A*^ mutant animals. The p.Q782*, p.A109P, and p. G201W variants failed to rescue lethality or reduced brain sizes (**Figure 2L-M**) consistent with them being severe loss-of-function variant alleles. However, p.L573V restored both viability and brain size (**Figure 2K, M**) in some *Ankle2*^*A*^ animals (**Figure 2M**), indicating that this variant is a mild hypomorphic allele.

### Ankle2 localizes to the ER and nuclear envelope and is required for their integrity

The tagged genomic rescue construct, *Ankle2-GFPR* (**Figure 3A-B**), as well as the endogenously tagged Ankle2^IGFP^ (**Figure S3**) show that *Ankle2* is expressed in most cells of the third instar larval brain. The protein appears to be localized to the cytoplasm of all cells including neuroblasts (arrows in **Figure 3A-B**). However, in a subset of cells, the protein is clearly enriched at the nuclear envelope (arrowhead). To determine the dynamics of Ankle2 protein localization, we performed live imaging. As shown in **Movies S1 and S2**, the protein is recruited to the nuclear envelope at the initiation of mitosis and remains associated with the nuclear envelope until briefly after cytokinesis.

**Figure 3:**
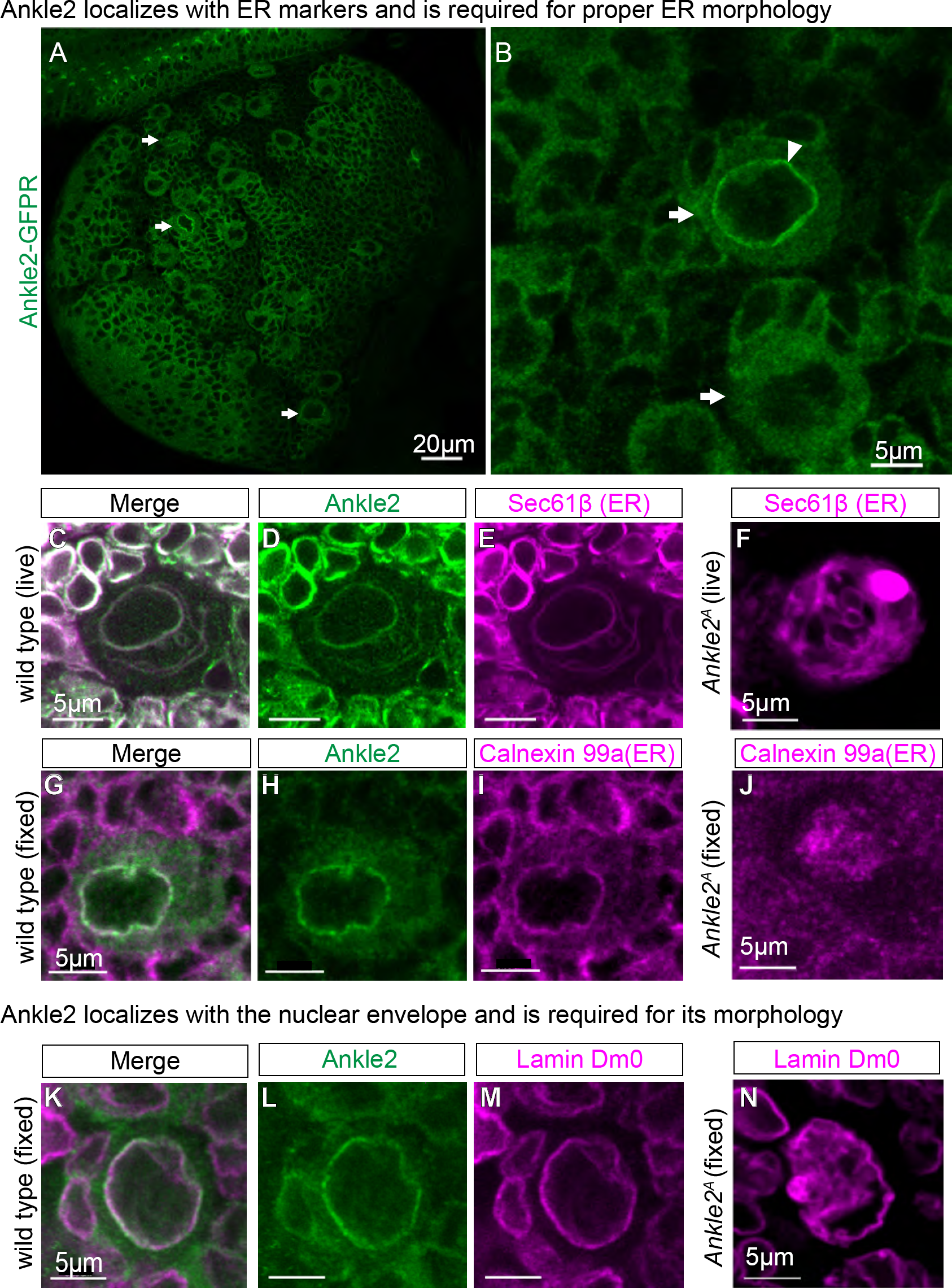
Ankle2 localizes to the ER and is dynamically expressed in the brain. (A-B) 3^rd^ instar larval brains from *Ankle2-GFPR* animals stained for GFP to document Ankle2 expression and localization in (A-B) single slices (arrows point to neuroblasts). (C-E) Live imaging of Ankle2 (D, green) and the ER labeled with Sec61B (E, *da-GAL4, UAS*-Sec61β-tdTomato, purple) (Summerville *et al.*, 2016) shows strong colocalization. (F) Live *Ankle2*^*A*^ mutant neuroblast display aberrant Sec61β-tdTomato expression. (G-I) Fixed *Ankle2-GFPR* animals highlighting Ankle2-GFP (green) and another ER marker (Calnexin 99a, purple) (Riedel *et al.*, 2016). (J) Fixed *Ankle2* mutant animals (*Ankle2*^*A*^) display aberrant ER structures (Calnexin 99a, purple). (K-M) Ankle2 (L, green) colocalizes with some portions of the nuclear envelope (Lamin Dm0, purple). (N) *Ankle2* mutant animals (*Ankle2*^*A*^) display disrupted nuclear envelope structure (Lamin DmO, purple). See also **Figure S3** and **Movie S1-S2**.

To determine precisely where Ankle2 is localized, we performed live imaging of brains from animals carrying *Ankle2-GFPR* and a transgene that labels the ER: *da-Gal4>UAS-Sec61β-tdTomato* (Summerville *et al.*, 2016). In neuroblasts (large cell in **Figure 3C-E**), the Ankle2 protein fully colocalizes with Sec61β at the nuclear envelope as well as the ER. In the surrounding neurons (small cells), much of the cytoplasm is colabeled. We also counter-stained fixed samples with Calnexin 99a, another ER marker (Riedel *et al.*, 2016). Again, Ankle2 localizes to the nuclear envelope and the ER, but in fixed samples, the ER structure is less obvious than in live imaging (**Figure 3G-I**).

To determine if Ankle2 is required for proper ER structure, we performed live imaging of *Ankle2*^*A*^ mutant neuroblasts expressing Sec61β-tdTomato (**Figure 3F**). When compared to wild type (**Figure 3E**), *Ankle2*^*A*^ mutants display highly aberrant Sec61β localization in many NBs (25°C). In addition, we stained fixed *Ankle2*^*A*^ mutant neuroblasts with Calnexin 99a and found that *Ankle2*^*A*^ mutants also display irregular Calnexin 99a localization (**Figure 3J** versus **3I**), suggesting that even a partial loss of Ankle2 disrupts ER and possibly nuclear envelope structure. Indeed, the morphology of the nuclear envelope is aberrant and convoluted in some *Ankle2*^*A*^ mutant cells when stained with Lamin DmO (Riemer *et al.*, 1995), a nuclear envelope marker (compare **Figure 3N** with **Figure 3K-M**). Hence, Ankle2 is required for proper ER and nuclear envelope morphology.

### *Ankle2* mutations affect the asymmetric localization of neuroblast determinants

Due to the reduced cell proliferation and reduced neuroblast number in *Ankle2*^*A*^ third instar brains (Yamamoto *et al.*, 2014), we sought to explore neuroblast division in more detail. Neuroblast polarity during division relies on the function of the highly conserved apically localized Par complex, which consists of Bazooka (Par3) (Schober *et al*., 1999), Par6 (Petronczki and Knoblich, 2001), and atypical Protein Kinase C (aPKC) (Rolls *et al.*, 2003). Once activated, the Par complex is responsible for restricting Miranda (Mira) and other cell fate determinants to the basal domain of neuroblasts. After division in most neuroblasts, the basal domain will become the ganglion mother cell, which divides again to produce neurons or glia (Betschinger *et al*., 2003; Atwood and Prehoda, 2009). Several proteins have been implicated in regulating the Par complex (Chabu and Doe, 2009; Andersen *et al.*, 2012; Bonaccorsi *et al.*, 2007; Atwood *et al.*, 2007), including those associated with cell cycle regulation (Chabu and Doe, 2008; Lee *et al.*, 2006; Wang *et al.*, 2007; Wang *et al.*, 2006).

Staining of third instar *Ankle2*^*A*^ brains with anti-Bazooka, Par6, aPKC, and Mira revealed severe localization defects of these proteins in greater than 40% of metaphase neuroblasts during asymmetric division (**Figure 4A-L**, quantified in **Figure 4M-P**) in both *Ankle2*^*A*^ and trans-heterozygous animals (*Ankle2*^*A*^/*Ankle2*^*CRIMIC*^). These defects are rescued by the genomic construct (**Figure 2D**), *Ankle2-GFPR* (**Figure 4D, H, L, M-P**). Finally, we performed live imaging of 3^rd^ instar larval brains of wild type (**Movie S3**) and *Ankle2*^*A*^ mutants labeled with Mira-RFP and Histone-GFP (**Movies S4-S6**). As shown in **Movies S4-S6**, neuroblasts exhibit abnormal Mira localization as well as some instances of failed division including DNA segregation defects, chromatin bridges, and cytokinesis defects (**Movies S4-S5**).

**Figure 4:**
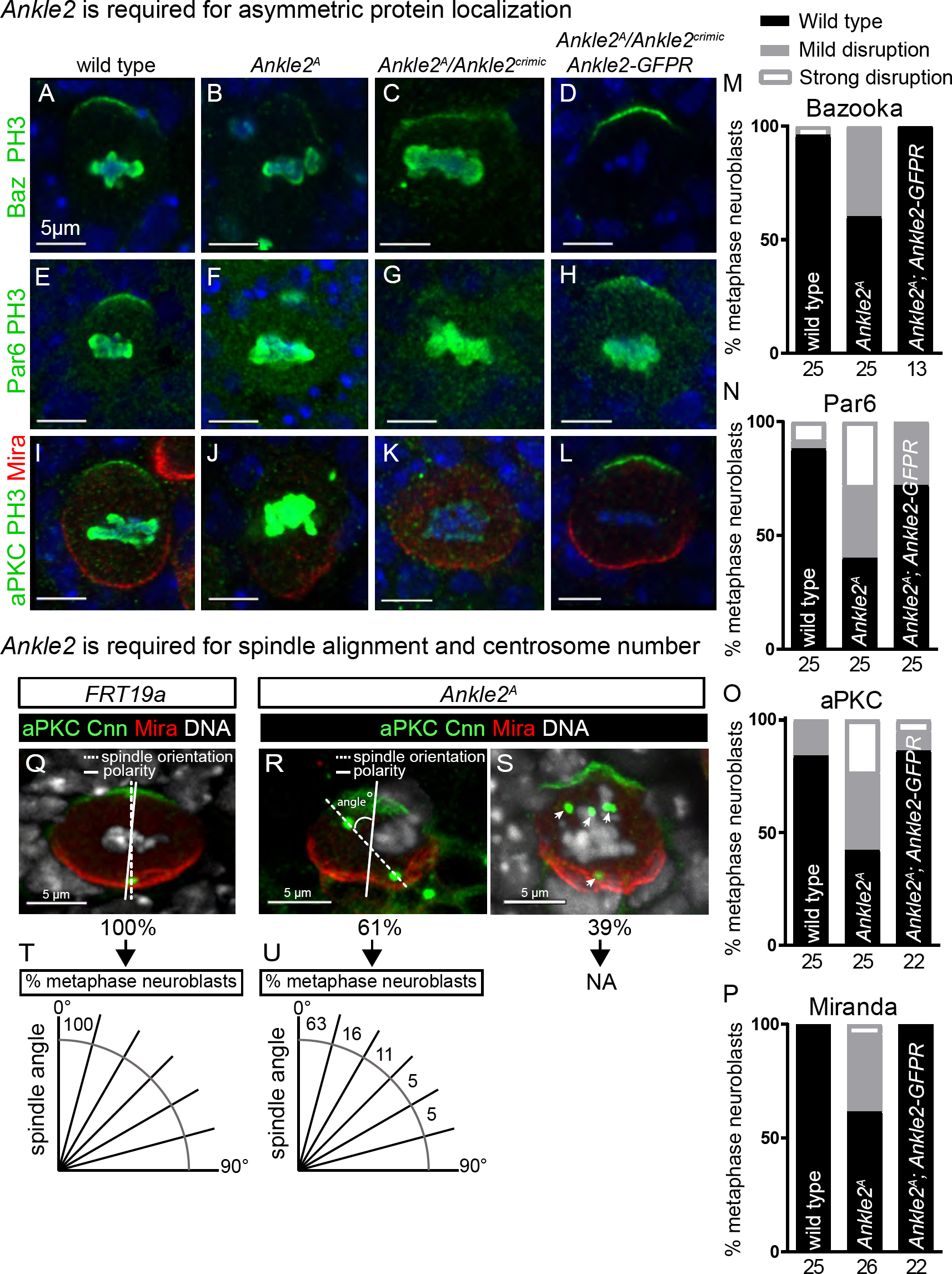
*Ankle2* mutations affect asymmetric division, spindle alignment and centrosomes. Metaphase neuroblasts stained with Baz (green, A-D), Par6 (green, E-H), aPKC (green, I-L) and Mira (red, I-L) are shown in wild type (A, E, I), *Ankle2*^*A*^/Y hemizygous (B, F, J), *Ankle2*^*A*^/*Ankle2*^*CRIMIC*^ transheterozygous (C, G, K) and rescued (D, H, L) animals. Phospho-Histone H3 (PH3, green, A-C, E-K) was used to identify cell cycle stage. * notes samples where PH3 was not used. (M-P) quantification of phenotype severity demonstrate that *Ankle2* is required for protein localization during asymmetric division in numerous cells. Below each graph, the # of neuroblasts counted for each genotype is noted. (Q-U) Metaphase neuroblasts stained with aPKC (green) and Mira (red) to mark the polarity axis and DNA (white) and CNN (green puncta) to highlight the spindle axis from (Q) wild type (*FRT19a*) and (R-S) *Ankle2* mutant neuroblasts. The angle between the spindle axis and polarity axis is measured and % of metaphase neuroblasts is plotted in 15° intervals and is shown in (T-U). See also **Movie S3-S6**.

For proper neuroblast division to occur, cells must not only asymmetrically localize Par complex members and cell fate determinants, they must also align the mitotic spindle so that divisions segregate cell fate determinants to the proper daughter cell (Cabernard and Doe, 2009). In wild type neuroblasts, the mitotic spindle is aligned parallel to the polarity axis (**Figure 4Q, T**). In some *Ankle2*^*A*^ mutant neuroblasts, we noted that spindle alignment appeared disrupted. To quantify these defects, we measured the axis of division using DNA and Centrosomin (CNN) (Lucas and Raff, 2007) to highlight centrosome placement relative to the localization of cell polarity proteins aPKC and Mira (**Figure 4Q-U**). Surprisingly, we found that nearly 40% of *Ankle2*^*A*^ mutant neuroblasts contained supernumerary centrosomes (**Figure 4S**). In the remaining 60% of *Ankle2*^*A*^ mutant metaphase neuroblasts with obvious aPKC/Mira localization, we found varying degrees of mitotic spindle alignment defects (compare **Figure 4Q-R** and **4T-U**), showing that Ankle2 is also required for proper spindle alignment in neuroblast division. Together, these results show that Ankle2 plays a prominent role in asymmetric protein localization, spindle alignment, and cell division of neuroblasts.

### Ankle2 interacts with VRK1/Ballchen

The *C. elegans* homologue of Ankle2, lem4L, was previously shown to physically and genetically interact with VRK1, the homologue of Ballchen (Ball) in flies (Asencio *et al.*, 2012). Lem4L and VRK1 in worms localize to the nuclear envelope of the 2-cell stage embryo (Asencio *et al.*, 2012). In contrast, Ball is localized to the nucleus in fly neuroblasts (Yakulov *et al.*, 2014). Interestingly, human *VRK1* pathogenic variants cause reduced brain size and microcephaly as well as axonal neuropathy in affected patients (Gonzaga-Jauregui *et al.*, 2013; Renbaum *et al.*, 2009). Hence, to characterize the relationship between Ankle2 and Ball/VRK1, we analyzed the expression and localization of Ball and Ankle2 during neuroblast cell division (**Figure 5A-D**). During interphase, Ankle2 and Ball do not colocalize as Ankle2 is in the cytoplasm and ER whereas Ball is in the nucleus (**Figure 5A**). During the mitotic prophase, Ankle2 accumulates at the nuclear envelope but the proteins do not seem to colocalize (**Figure 5B**). However, upon fragmentation of the nuclear envelope during metaphase, Ball is briefly present throughout the cytoplasm (**Figure 5C**). Yet, during telophase, Ball is quickly recruited back to the nucleus and briefly enriched at the nuclear envelope (**Figure 5D**; **Movie S7**). Interestingly, the spatial restriction of Ball in *Ankle2*^*A*^ mutants during interphase is abolished in many neuroblasts as Ball localizes throughout the cell, a phenotype that is not observed in wild type brains (**Figure 5E-F).** In summary, Ankle2 is required for proper nuclear localization of Ball in *Drosophila*.

**Figure 5:**
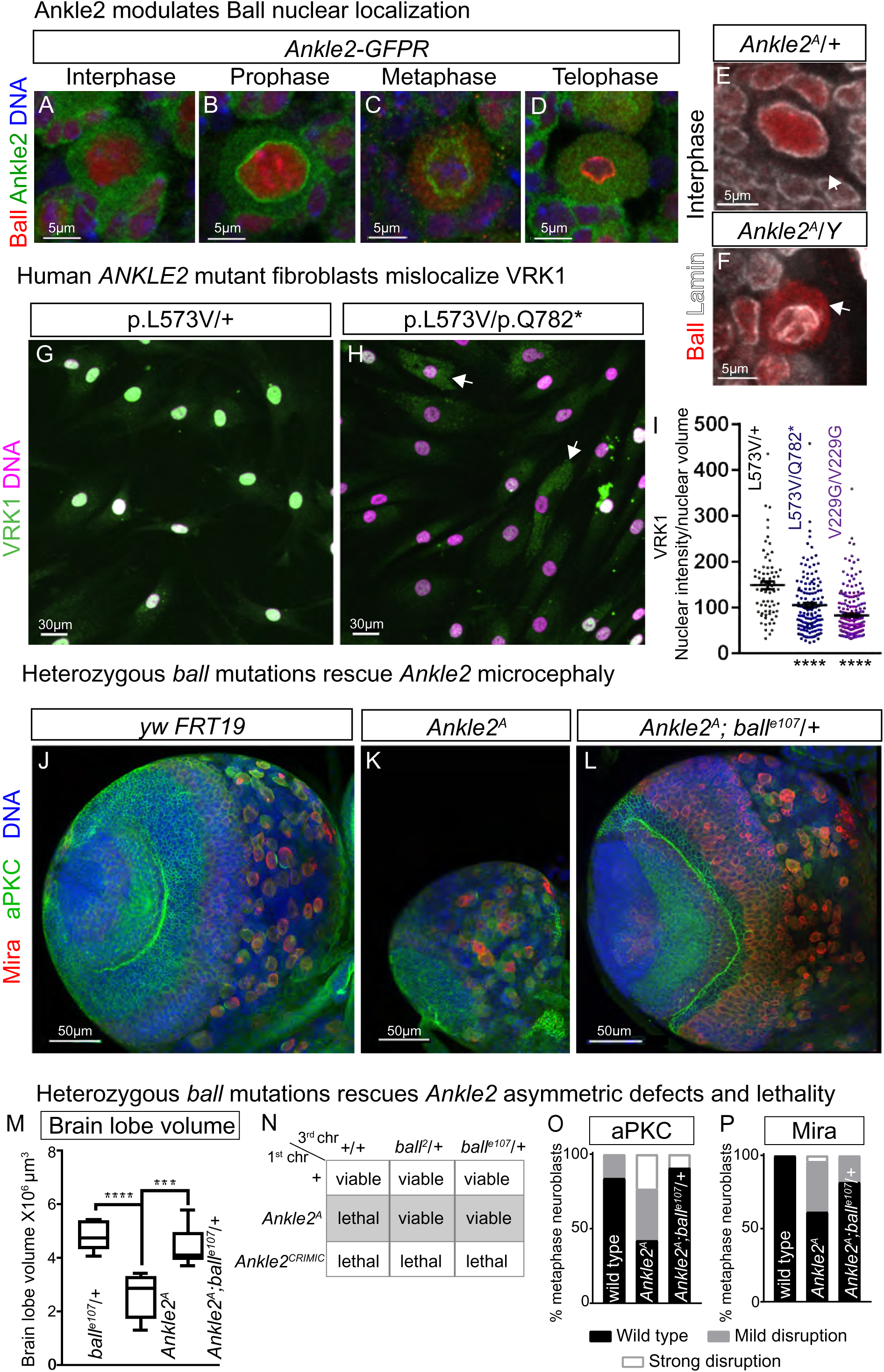
Ankle2 controls Ballchen/VRK1 localization and function. (A-D) Immunostaining of Ankle2-GFPR (green) and Ball (red) show dynamic localizations during the cell cycle. (B) Ankle2 is localized at the nuclear envelope at prophase. Ball is nuclear (A-B) until nuclear envelope breakdown (C) and then localizes to the cytoplasm through the end of mitosis (D). (E-F) Immunostaining of Ball (red) and Lamin (white) in *Ankle2*^*A*^*/+* heterozygotes and *Ankle2*^*A*^*/Y* hemizygous mutants. Note that Lamin is disrupted and Ball becomes mislocalized throughout the cytoplasm during interphase in *Ankle2*^*A*^*/Y* hemizygous mutants. (G-H) Confocal projections of immunostaining of VRK1 (green) and DNA (purple) in primary human fibroblasts from (G) parental unaffected (p.L573V/+) and (H) an ANKLE2 compound heterozygous patient (p.L573V/p.Q782*). VRK1 is mislocalized in fibroblasts carrying microcephaly associated *ANKLE2* variants (p.L573V/p.Q782* and p.V229G/p.V229G), quantified as nuclear intensity in (I). Arrows in (H) indicate cytoplasmic VRK1 staining, which is minimal in (G) control fibroblasts. (J-L) Partial projections of 3^rd^ instar larval brains stained with apical marker aPKC (green) and basal marker Miranda (red) in (J) wild type (*y,w, FRT19a*), (K) *Ankle2*^*A*^, and (L) *Ankle2*^*A*^;;*ball*^*e107*^/+ animals. Note that removal of a single copy of *ball* rescues the phenotypes of *Ankle2*^*A*^. (M) Quantification of 3^rd^ instar larval brain size (as shown in (J-L)). N≥6. One-way ANOVA with multi-comparison post test. ****p<0.0001, ***p<0.001. Box plots hinges represent the 25^th^ to 75^th^ percentiles, a line is at the median, and whiskers represent min to max. (N) *Ankle2*^*A*^, but not *Ankle2*^*CRIMIC*^, lethality is rescued with introduction of multiple *ball* heterozygous mutations. (O-P) Quantification of (O) aPKC or (P) Mira crescent intensity in 3^rd^ instar metaphase neuroblasts in wild type (*y w FRT19a*), *Ankle2*^*A*^ and rescued *Ankle2*^*A*^; *ball*^*e107*^*/+* animals, demonstrating that *Ankle2* asymmetric division phenotypes are rescued with *ball* heterozygosity. Note that wild type and *Ankle2*^*A*^ quantifications were shown in Fig 3. See also **Figure S4-S6 and Movie S7.**

To determine whether ANKLE2 regulates Ball/VRK1 subcellular localization in human cells, we assayed VRK1 localization in human fibroblasts. In reference human primary fibroblasts (parental variant p.L573V/+), VRK1 is localized to the nucleus (**Figure 5G**). However, fibroblasts from microcephaly patients carrying compound heterozygous variants in *ANKLE2* (p.L573V/p.Q782* and p.V229G/p.V229G) display significantly reduced VRK1 intensity in the nucleus (**Figure 5H**, quantified in **Figure 5I**) and increased cytoplasmic staining in non-dividing cells (arrows in **Figure 5H**) with no significant change in overall VRK1 intensity (**Figure S4**). These data argue for a conserved role between fruit flies and human for ANKLE2 in restricting VRK1 to the nucleus.

Given that Ankle2 is required to maintain Ball/VRK1 in the nucleus during interphase, it is possible that Ball/VRK1 is ectopically active in the cytoplasm of *Ankle2*^*A*^ mutants and inhibits or promotes phosphorylation of proteins not normally encountered in the biological homeostatic state. Reducing the level of Ball/VRK1 may therefore alleviate the phenotype associated with the reduction in Ankle2 protein. Indeed, we observe evidence for strong dominant interactions between *Ankle2*^*A*^ and *ball* (multiple alleles). *Ankle2*^*A*^ animals are pupal lethal and have reduced brain volumes (compare **Figure 5J** to **5K**). However, removal of one copy of *ball*, akin to a heterozygous deletion CNV resulting in haploinsufficiency in human, restores brain development (**Figure 5L-M**) and suppresses the lethality of *Ankle2*^*A*^ mutants (**Figure 5N**). Importantly, loss of one copy of *ball* (*ball*^*e107*^) in *Ankle2*^*A*^ mutants also restores the asymmetric protein localization of aPKC and Mira crescents in metaphase neuroblasts (**Figure 5O-P**). Hence, a partial reduction of Ball/VRK1 activity rescues *Ankle2*^*A*^ mutants, providing strong evidence for a gene dosage sensitive locus. However, removing both copies of *ball* in wild type animals leads to pupal lethality (Cullen *et al.*, 2005), causes severely reduced brain volumes in 3^rd^ instar larvae (Herzig *et al.*, 2014), and does not rescue *Ankle2*^*A*^ animals, emphasizing that the gene dosage and balance of the protein levels is critical. Indeed, a severe loss of function allele, *Ankle2*^*CRIMIC*^, cannot be suppressed by reducing Ball/VRK1 activity (**Figure 5N**). In summary, these data demonstrate that both Ankle2 and Ball/VRK1 control the distribution of asymmetric determinants, and experimental evidence reveals an antagonistic relationship between both proteins.

### The Ankle2-Ball pathway modulates aPKC and L(2)gl

Due to the similarities in defects observed with loss of *Ankle 2* or *aPKC*, including mislocalization of Par6 and Mira (Kim *et al.*, 2009), decreased cell divisions, and reduced neuroblast clone volume (Rolls *et al.*, 2003), we hypothesized that the activity of aPKC, an important mediator of neuroblast asymmetric division (**Figure 6A**), might be affected. aPKC phosphorylation (Kim *et al.*, 2009) or abundance could be modulated by Ankle2. We therefore assessed both total and phosphorylated aPKC levels in third instar larval brains using an antibody specific for human p-aPKC T410 (T422 in flies). This phosphorylation site is located in its activation loop and was shown to be important for its kinase activity (Kim *et al.*, 2009). Phosphorylation of aPKC (T422) relative to total aPKC is decreased in *Ankle2* mutants (**Figure 6B**) and is restored with either addition of Ankle2-GFPR or reduction of *ball* (**Figure 6B**), consistent with the data presented in **Figure 5**. However, overexpression of aPKC or constitutively active aPKC (aPKC^ΔN^) (Betschinger *et al*., 2003) in *Ankle2*^*A*^ mutants did not rescue brain size or viability (data not shown).

**Figure 6:**
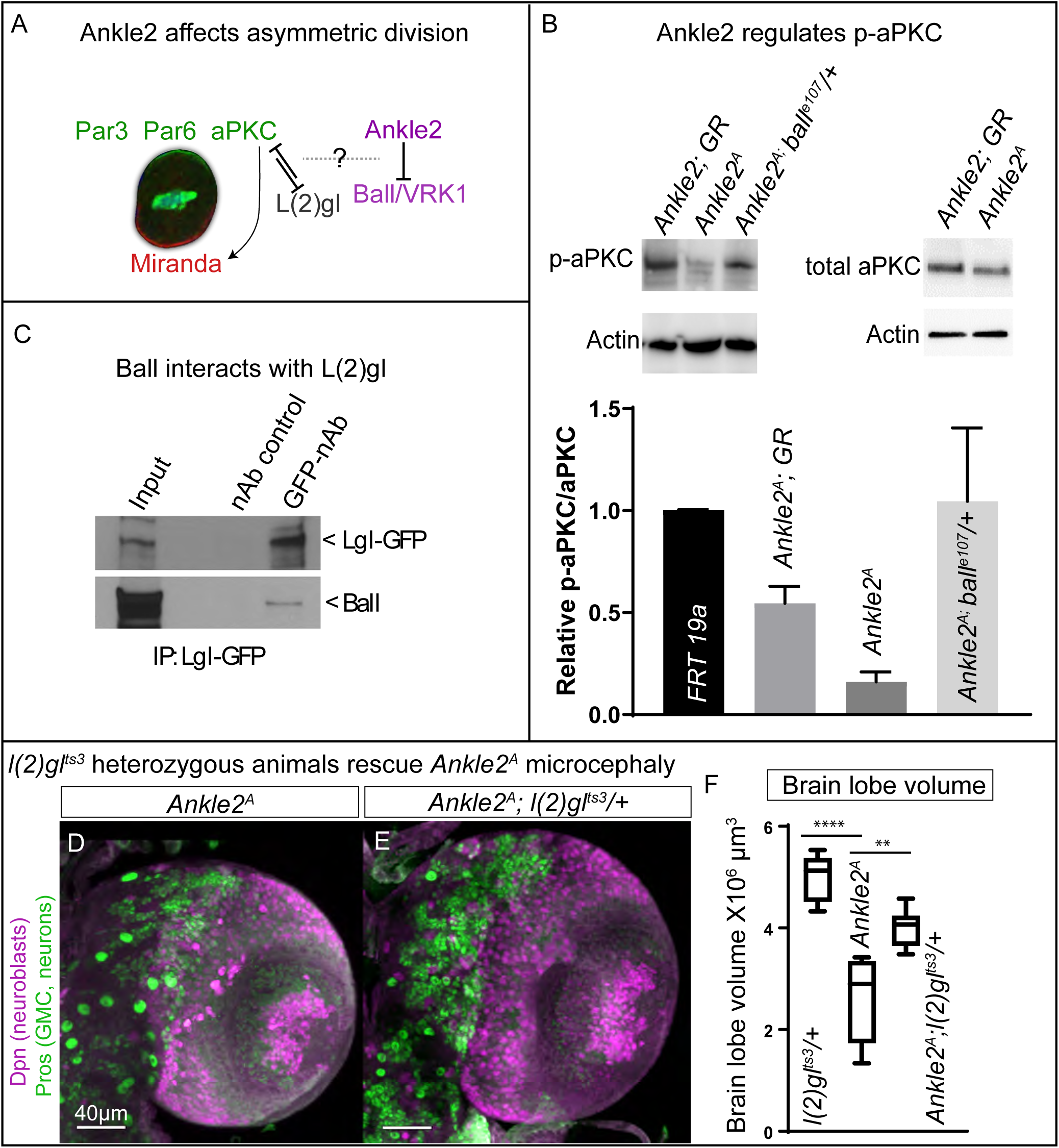
Ankle2 affects asymmetric division through aPKC and L(2)gl. (A) Ankle2 and Ball regulate asymmetric division. (B) Western analysis of phosphorylated aPKC in larval brains (mammalian T410 corresponds to T422 in *Drosophila*) from genomic rescue (*Ankle2*^*A*^; *Ankle2-GFPR*), *Ankle2*^*A*^, and *Ankle2*^*A*^*;; ball*^*e107*^*/+* animals or total aPKC levels in genomic rescue (*Ankle2*^*A*^; *Ankle2-GFPR*) and *Ankle2*^*A*^ *mutants*. Note that p-aPKC is reduced in *Ankle2* mutants but restored with introduction of genomic rescue (GR) or reduction of Ball and is quantified as the ratio of p-aPKC to total aPKC. N=3 replicates. (C) *In vivo* immunoprecipitation of L(2)gl-GFP using GFP-nAb in *L(2)gl*^*MI07575-GFSTF*^ larvae demonstrates L(2)gl can interact with Ball *in vivo*. (D,F) Partial projections of third instar larval brains stained for Dpn (purple, neuroblasts) and Pros (green, daughter cells and neurons) of (D) *Ankle2*^*A*^ and (E) *Ankle2*^*A*^*/+;l(2)gl*^*ts3*^ mutant animals raised at 22°C with brain volume quantified in (F). Note that reduction of L(2)gl in an *Ankle2*^*A*^ hemizygous animals rescues brain size defects and lethality at 22°C. Box plots hinges represent the 25^th^ to 75^th^ percentiles, a line is at the median, and whiskers represent min to max. See also **Figure S7**.

aPKC has been shown to physically interact with L(2)gl (Betschinger *et al*., 2003), a regulator of apico-basal polarity that inhibits the function of aPKC (Atwood and Prehoda, 2009; Wirtz-Peitz *et al*., 2008). *aPKC* and *l(2)gl* genetically interact as removal of one copy of *aPKC* suppresses *l(2)gl* loss of function phenotypes (Rolls *et al.*, 2003), and aPKC has been shown to phosphorylate L(2)gl to control its plasma membrane or cortical release (Betschinger *et al*., 2003). When aPKC is active, L(2)gl is phosphorylated and released from the cortex; once released, it no longer binds to aPKC or inhibits its function. Because aPKC and L(2)gl interact, the Ankle2-Ball pathway may affect L(2)gl. We therefore assessed whether L(2)gl physically interacts with the Ankle2-Ball pathway using immunoprecipitation of a GFP-tagged L(2)gl from third instar larval brains and found that Ball indeed interacts with L(2)gl (**Figure 6C**).

The reduced aPKC activity that we observe may be associated with a gain of function of L(2)gl. Therefore, to determine whether removal of one copy of *l(2)gl* suppresses *Ankle2* associated phenotypes, we introduced a temperature sensitive mutation of *l(2)gl (l(2)gl*^*ts3*^) into the *Ankle2*^A^ mutant background and found that reducing *l(2)gl* in *Ankle2*^*A*^ mutants at 22°C (**Figure 6D-F**) and 25°C (not shown) indeed partially restored brain size. *Ankle2*^*A*^ is pupal lethal at 22°C, but when combined with a heterozygous *l(2)gl* mutant allele, some *Ankle2*^*A*^ animals survive to adulthood. However, unlike the removal of one copy of *ball*, these animals die a few days after eclosion. In summary, Ankle2 and Ball interact with the apical-basal polarity regulators aPKC and L(2)gl (**Figure 6A**) and affect aPKC and L(2)gl activity by disturbing the asymmetric segregation of apical-basal polarity factors in neuroblasts.

### Disease associated variants in *VRK1* and its paralogs

Ten families have been described with biallelic variants in *VRK1* that cause a spectrum of neurologic diseases including 6 individuals with microcephaly (Feng *et al.*, 2018; Gonzaga-Jauregui *et al*., 2013; Najmabadi *et al*., 2011; Nguyen *et al.*, 2015; Renbaum *et al.*, 2009; Shaheen *et al*., 2018; Stoll *et al.*, 2016) (**Table S1; Figure S5**). The family structures suggest either a sporadic or recessive neurological disease trait; historical consanguinity in 3/10 pedigrees implicate an autosomal recessive locus. Screening the BHCMG and BG databases identified two additional families with potentially biallelic variants in *VRK1*. (**Table S2, Figure S5**). These cases suggest that like *ANKLE2* **(Figure 1C and D)**, a heterogenous set of variant alleles in *VRK1* are associated with neurologic disease and microcephaly.

It was previously shown that fly genes with more than one human homolog, especially those that are evolutionarily conserved, have an enriched association with OMIM disease phenotypes (Yamamoto *et al*., 2014). We searched the BHCMG database to establish if damaging variants in paralogs of *VRK1* are associated with disease. Predicted deleterious, biallelic variants were found in two paralogs of *VRK1*: *VRK2* is associated with very small eyes and *VRK3* with severe microcephaly (**Table S2, Figure S6**).

### NS4A targets the Ankle2 pathway

*Drosophila* has been developed as a model of viral infection (Harsh *et al.*, 2018; Liu *et al.*, 2018), and we recently showed that expression of the Zika protein NS4A results in reduced brain size in *Drosophila*. Strikingly, NS4A expression in *Ankle2*^*A*^/+ heterozygous animals leads to a more severe phenotype than NS4A expression in a wild type background, and these animals display brain phenotypes that mimic *Ankle2*^*CRIMIC*^ null mutants (Shah *et al.*, 2018). These data again suggest that levels of Ankle2 protein are critical, and hence, expression of NS4A in neuroblasts may cause microcephaly by affecting aPKC and Miranda localization. Indeed, expression of NS4A in neuroblasts (*insc-GAL4>UAS-NS4A*) affects the apical aPKC localization and leads to an expansion of the Mira domain at the basal membrane (**Figure 7A-E**). In the metaphase neuroblasts that express NS4A, we also note spindle orientation defects in some cells (**Figure 7G-H**), similar to *Ankle2*^*A*^ animals shown in **Figure 4**. Hence NS4A targets the Ankle2 pathway; this is further strengthened with the observation that when NS4A is expressed in neuroblasts of *ball* heterozygous animals, aPKC and Mira crescents are restored to their wild type patterns (**Figure 7D-F**) and spindle orientation defects are rescued (compare **Figure 7I** with **G** and **H**). In summary, the Zika virus protein NS4A targets the Ankle2 pathway and affects asymmetric distribution of cell fate determinants, leading to defects in neuroblast division.

**Figure 7:**
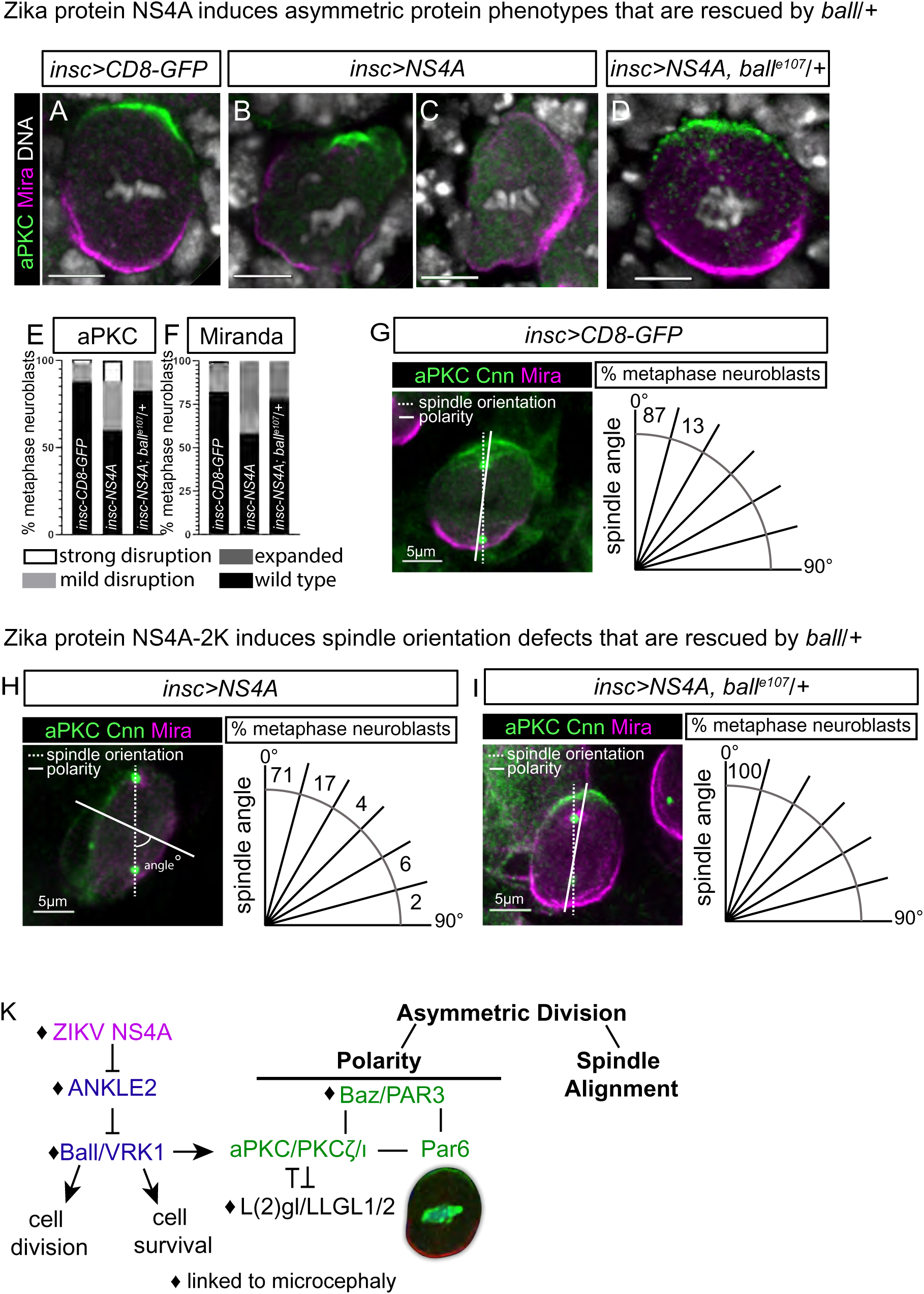
Zika NS4A targets the Ankle2 pathway. 3^rd^ instar metaphase neuroblasts stained for aPKC (green) and Mira (purple) with neuroblast specific (*insc-GAL4*) expression of (A) CD8-GFP, (B-C) NS4A, and (D) NS4A in *ball*^*e107*^ heterozygous animals. aPKC and Mira crescent intensities are quantified in (E-F). (G-I) aPKC (green), CNN (green), and Mira (purple) staining of metaphase neuroblasts with *insc-GAL4* expression of (G) CD8-GFP, (H) NS4A, and (I) NS4A in *ball*^*e107*^ heterozygous animals. The angle between the spindle axis and polarity axis is measured and % of total metaphase neuroblasts is plotted at 15° intervals. Expression of NS4A causes localization defects of aPKC and Mira and spindle orientation defects in metaphase neuroblasts. (K) ZIKV NS4A inhibits the ANKLE2/VRK1 pathway, which regulates asymmetric determinant localizations as well as the division axes.

## DISCUSSION

We investigated the biological basis for *ANKLE2* associated microcephaly. We report six new patients with microcephaly that carry mutations in *ANKLE2* and show that three variants identified in probands cause a loss of *ANKLE2* function when tested in flies (**Figure 1 and 2**), providing compelling evidence that its loss causes reduced brain size in flies and severe microcephaly (Z-score < −2.5) in humans. *Ankle2* is a dosage sensitive locus whose product is inhibited by the Zika virus protein NS4A. We show that Ankle2, like NS4A (Shah *et al.*, 2018), is localized to the ER, and that it targets the nuclear envelope during mitosis. Loss of *Ankle2* affects the nuclear envelope and ER distribution and results in a redistribution of Ball/VRK1, a kinase that is normally localized to the nucleus except when the nuclear envelope breaks down during mitosis (**Figure 5**). Loss of *Ankle2* disrupts the localization of neuroblast apical-basal polarity determinants such as aPKC, Par 6, Baz, and Miranda, and aPKC phosphorylation is reduced by *Ankle2* mutations. Importantly, loss of one copy of *ball* or *l(2)gl* suppresses the reduced brain volume associated with a partial loss of *Ankle2*, suggesting that much of the biological function of Ankle2 is modulated by aPKC and L(2)gl. Finally, the negative effect of NS4A on the activity of ANKLE2 can also be suppressed by removal of one copy of *ball*, suggesting the following pathway: NS4A ┤ ANKLE2 ┤ Ball/VRK1 → L(2)gl/LLGL1 ┤ aPKC. This represents an important pathway regulated by ANKLE2 that we show plays an important role in neuroblast stem cell divisions in flies and microcephaly phenotypes in humans.

Interestingly, the above pathway links environmental cues with several genetic causes of sporadic and AR microcephaly in human; moreover, it implicates this pathway in microcephaly accompanying congenital infection. As one example of the latter, the Zika virus has been shown to cross the infant Blood Brain Barrier (Mlakar *et al.*, 2016) and has been identified in radial glial cells (Li *et al.*, 2016), as well as intermediate progenitor 7 cells and neurons (Lin *et al.*, 2017). We propose that NS4A affects the function of Ankle2 leading to the release of Ball/VRK1 from the nucleus. We speculate that this in turn affects the phosphorylation of aPKC and L(2)gl directly by masking phosphorylation sites or indirectly by promoting the activity of one or more phosphatases. Loss of VRK1 has been shown to cause microcephaly and some variant alleles are also associated with pontocerebellar hypoplasia (PCH) in humans (Gonzaga-Jauregui *et al.*, 2013; Renbaum *et al.*, 2009), consistent with the loss of *ball* in flies that causes a severe reduction in brain size (Yakulov *et al.*, 2014). Note that ANKLE2, VRK1, LLGL1, and aPKC, as well as other components of the apical complex like PARD3 are all present in radial glial cells during cortical development (Ayoub *et al.*, 2011). These data suggest that ANKLE2 and its partners such as LLGL1 and asymmetric determinants are important proteins during neural cell proliferation and that the proper levels and relative amounts of these proteins determine how many neurons will eventually be formed in vertebrates. Our data also indicate variant alleles at either *ANKLE* or *VRK1* are responsible for some causes of embryonic lethality and severe congenital microcephaly.

*LLGL1* has recently been shown to play an important role in radial glia in mice during neurogenesis, and its loss in clones increases the number of divisions (Beattie *et al.*, 2017). In addition, aPKCζ/λ localizes at the apical membrane of proliferating neural stem cells in chicken embryos during division and has been shown to provide an instructive signal for apical assembly of adherens junctions (Ghosh *et al.*, 2008). Mouse knockouts of aPKCλ (Soloff *et al.*, 2004) and aPKCζ (Seidl *et al.*, 2013) are embryonic lethal; however aPKCζ knockouts are viable (Leitges *et al.*, 2001), perhaps suggesting redundant functions within the atypical PKC family. These proteins have not been linked to microcephaly in mice, but conditional removal of an apical complex protein Pals1 in cortical progenitors resulted in complete cortex loss (Kim *et al.*, 2010). Finally, Numb is asymmetrically localized by the Par complex protein in *Drosophila*, segregated to the daughter cell during asymmetric cell division (Wirtz-Peitz *et al*., 2008), and essential for daughter cells to adopt distinct fates (Bhalerao *et al.*, 2005). In mice, Numb localization is also asymmetric and null mutations exhibit embryonic lethality, neural tube closure defects, and premature neuron development (Zhong *et al.*, 2000). These data indicate that asymmetric division may be important for vertebrate neuronal development, but microcephaly is not a phenotype that typically associates with loss of the mice homologues of asymmetric localized determinants studied *Drosophila*. However, the observations reported here indicate that the ANKLE2/PAR complex pathway is evolutionarily conserved from flies to human, although the precise mechanisms remain to be determined as different cells may use this pathway in different contexts (Suzuki and Ohno, 2006).

In order to determine whether predicted deleterious biallelic variants in PAR complex encoding genes or their paralogs associated with a neurologic disease trait, we searched the BHCMG database for mutations associated with neurological disease. We found homozygous predicted deleterious missense variants in *PARD3B* (c.1222G>A, p.G408S) in a patient that has microcephaly (**Table S2, Figure S7**) and compound heterozygous mutations in *PARD3B* (c.1654G>A;p.A552T) that are associated with other neurological defects (**Table S2, Figure S7**). The human orthologue of L(2)gl, *LLGL1*, is deleted in Smith-Magenis syndrome (SMS) (Smith *et al.*, 1986) and 86-89% of the SMS patients have brachycephaly (Greenberg *et al.*, 1996). These observations extend the mutational load beyond *ANKLE2* and *VRK1* and suggest an association between congenital disease and variants within the PAR complex (**Table S2, Figure S7**).

Aurora A (AurA) kinase has been shown to phosphorylate the Par complex (Wirtz-Peitz *et al*., 2008) as well as L(2)gl (Carvalho *et al.*, 2015) and regulates cortical polarity and spindle orientation in neuroblasts (Lee *et al.*, 2006). The aberrant localization of Ball/VRK1 in *Ankle2* mutants may lead to gain of function phenotypes that are highly dosage sensitive, as they can be repressed by removing a single copy of Ball/VRK1 in *Ankle2*^*A*^. Mislocalized Ball/VRK1 may mask or interfere with the function of AurA in neuroblast asymmetric division as they share similar kinase substrate consensus sequences (Sanz-García *et al.*, 2011; Ferrari *et al.*, 2005). Future studies are needed to assess Ball/VRK1 redundancy or interference with AurA function.

Another possible evolutionarily parallel with implications in multicellular organismal development is the genetic interaction between the *C. elegans* homologue of VRK1 and an ANKLE2-like protein at the two cell stage (Asencio *et al.*, 2012). Whereas VRK1 in both *Drosophila* and humans (Nichols and Traktman, 2004) is localized to the nucleus, except during mitosis when the nuclear envelope is broken down (**Figure 5**), the worm VRK1 protein is localized to the nuclear envelope. The worm ANKLE2-like protein, Lem-4L, also interacts with the phosphatase PP2A (Asencio *et al.*, 2012), and Chabu and Doe (2009) noted that the fly PP2A regulates neuroblast asymmetric division by interacting with aPKC and excluding it from the basal cortex (Chabu and Doe, 2009). PP2A also antagonizes the phosphorylation of Baz by PAR-1 to control apical-basal polarity in dividing embryonic neuroblasts (Krahn *et al*., 2009) and regulates Baz localization in other cells such as neurons (Nam *et al*., 2007). This raises the possibility that the Ankle2 pathway also acts with PP2A in neuroblast asymmetric division.

Here, we identified a novel pathway that plays a significant role in neuroblast asymmetric division. By combining functional studies in *Drosophila* together with human subject data, we have linked several microcephaly-associated genes and congenital infection to a single genetic pathway. These studies allowed us to highlight conserved functions of the ANKLE2 pathway, and provide mechanistic insight on how a Zika infection might affect asymmetric division. This *ANKLE2*-*VRK1* gene dosage sensitive pathway can be perturbed by genetic variants that disturb biological homeostasis resulting in neurological disease traits or by environmental insults such as Zika virus impinging on neurodevelopment. Hence, lessons learned from the study of rare diseases such as MCPH16/*ANKLE2* can provide insights into more common disease and potential gene by environmental interactions.

## Acknowledgements

We would like to thank many members of the Bellen lab for suggestions, and Karen Schulze for critical reading. We thank Chris Doe, Jurgen Knoblich, Jim Skeath, Kenneth Prehoda, Alf Herzig, Hiroyuki Ohkura, and the Bloomington Drosophila Stock Center for providing stocks and reagents, and the IDDRC Microscopy Core (NIH/NICHD U54 HD083092) for valuable input. This work was supported by NIH/NINDS F32NS092270 to N.L., Howard Hughes Medical Institute (HHMI) Medical Research Fellowship to A.J., the NIH/NINDS K08NS092898 and Jordan’s Guardian Angels to G.M.M., a jointly funded NHGRI and NHLBI grant to the Baylor-Hopkins Center for Mendelian Genomics (<u>UM1 HG006542</u>) and NIH/NINDS R35NS105078 to J.R.L, NIH U54NS093793, NIH R24OD022005, and the Huffington Foundation to H.J.B.. N.L. and H.C. are supported by HHMI, and H.J.B. is an Investigator of the Howard Hughes Medical Institute.

## Author Contributions

N.L. and H.J.B. conceived the project and designed experiments, and wrote and revised the manuscript with J.R.L. H.C. performed *in vivo* Ankle2 immunoprecipitations and assessed phosphorylation/total protein levels. A.J. assisted with brain volume measurements. A.J., M.W. and J.R.L performed primary fibroblast experiments, and human mutation studies. H.A., B.B.G., T.T., S.I., B.T., G.M.M., G.H.M., A.X.J., and R.D.C. ascertained clinical and molecular data of children with variants. B.T., B.A., P.S., and N.K. assisted with ZIKV experiments. N.L. performed all other experiments.

## Methods

### CONTACT FOR REAGENT AND RESOURCE SHARING

Further information and requests for resources and reagents should be directed to and will be fulfilled by the Lead Contact, Hugo J. Bellen.

### EXPERIMENTAL MODEL AND SUBJECT DETAILS

#### Drosophila melanogaster

The following fly lines were used: *FRT19a* (Yamamoto *et al.*, 2014), *Ankle2*^*A*^ (Yamamoto *et al.*, 2014), *Ankle2*^*CRIMIC*^ (this study), *Ankle2*^*IGFP*^(this study), *Ankle2-GFPR* (this study), *P{UASt-hANKLE2}VK37* (Yamamoto *et al.*, 2014), *P{UASt-hANKLE2 p.L537V}VK37* (this study), *P{UASt-hANKLE2 p.Q782*}VK37* (this study), *P*{*UASt-hANKLE2 p.A109P}VK37* (this study), *P*{*UASt-hANKLE2 p.G201W}VK37* (this study), *P{20XUAS-tdTomato-Sec61beta}attP2* (Summerville *et al.*, 2016), *ball*^*e107*^(Cullen *et al.*, 2005), *ball*^*2*^ (Herzig *et al.*, 2014), *l(2)glMI07575-GFSTF.0* (Nagarkar-Jaiswal *et al.*, 2015a), *l(2)gl*^*ts3*^*cn*^*1*^*sp*^*13*^ (Manfruelli *et al.*, 1996), *P*{*UASt-NS4Aug*} (Shah *et al.*, 2018), *P*{*UASt-CD8-GFP}* (Lee and Luo, 2001), *Actin-GAL4* (*P{Act5C-GAL4}17bFO1*) (Ito *et al.*, 1997), *inscuteable-GAL4* (*P{w[+mW.hs]=GawB}insc[Mz1407]*) (Luo *et al.*, 1994), *daughterless-GAL4 (P{w[+mW.hs]=GAL4-da.G32}UH1)* (Wodarz *et al.*, 1995), *ball-GFP (fTRG-823)* (Sarov *et al.*, 2016), *wor-mira-cherry P{w[+mC]=wor.GAL4.A}2,P{w[+mC]=UAS-mira.cherry}2/CyO)* (Cabernard and Doe, 2009), *P{His2Av[T:Avic\GFP-S65T]}62A* (Clarkson and Saint, 1999), *P{w[+mC]=UAS-aurA.Exel}2, M{ UAS-aPKC.ORF.3xHA}ZH-86Fb* (Bischof *et al*., 2013), *P{w[+mC]=UAS-aPKC.DeltaN}3* (Drier *et al.*, 2002). All flies were maintained at 22°C and grown on standard cornmeal and molasses medium in plastic vials. Crosses were performed at temperature indicated (18°C, 22°C, 25°C, or 29°C). Hemizygous males were analyzed as *Ankle2* mutants (which is on the X chromosome) and females were used for *Ankle2* heterozygous studies. All other studies contained males and females. Brain volume measurements were conducted in late 3^rd^ instar larvae (gut clearance, extruding spiracles). *Act-GAL4* was use to ubiquitously express ZIKV NS4A and Sec61β, *da-GAL4* was used to express human ANKLE2 constructs and aPKC, *insc-GAL4* was used to express NS4A in neuroblasts.

#### Human studies

All study subjects enrolled into the Baylor-Hopkins Center for Mendelian Genomics (BHCMG) provided informed consent for exome sequencing and study participation under the Baylor College of Medicine Institutional Review Board-approved protocol H-29697. BAB701 provided informed consent for molecular and genomic analysis under the Baylor College of Medicine Institutional Review Board-approved protocol H-9170. All study subjects enrolled through Baylor Genetics Laboratory (BGL) were analyzed on a retrospective basis, and only de-identified information is provided under the Baylor College of Medicine Institutional Board approved protocol H-41191. Patients were ascertained from the 7148 sequenced individuals in BHCMG or the ∼12500 sequenced individuals in the BGL by searching for biallelic variants with CADD scores >15 in conjunction with phenotypes of interest. Six male and two female patients were ascertained from the BHCMG database. The age of the patient is known for 3 individuals (2y, 7y, 32y). Two female and three male patients were ascertained from the BGL database. Ages at referral were 2 months, 7y, 12y, 20y, and 41y.

#### Human primary cultures

Fibroblasts were cultured in flasks containing Gibco DMEM (1x) with 4.5g/L D-Glucose, L-Glutamine, 25mM HEPES, HyClone FBS (10%), and Gibco Anti-Anti (1%) at 37 degrees Celsius with 5% CO2. Cultures of p.L573V/+ and p.L573V/p.Q782* were male; p.V229G/p.V229G was female.

### METHOD DETAILS

#### Generation of Ankle2 mutations and constructs

To generate *Ankle2*^*CRIMIC*^ by CRISPR-Cas9, two guide RNAs targeting *Ankle2*, 5’-ATAAAGTATTTTCTTAACGG<u>TGG</u>-3’ and 5’-TAATAATTTTAAATTCTCAT<u>TGG</u>-3’ with PAM sites underlined, were cloned into pCDF3 (Port *et al.*, 2014). Regions of homology targeting the 4^th^ coding intron were cloned into PM14 (Lee *et al.*, 2018) using 5’-ccatagctatggGCAATTCCTCAATGTCGAATTTACTGCTCA-3’ and 5’-ttatgcatATTTTCTTAACGGTGGGAAATTATAC-3’ to amplify the left arm for BstXI/NsiI cloning and 5’-tagcatgcATACTTTATTATTGCATTTGTTATAAGTATGAGA-3’ and 5’-tactcgagGCAAAGTTCCAGACCGTTTCTGATTTATC-3’ to amplify the right arm for conventional cloning with SphI/XhoI. This donor construct and two guide RNA constructs were injected into, *w;attP40(y+){nos-Cas9(v+)}/CyO* (Kondo and Ueda, 2013) embryos, and positive expression of *3XP3-GFP* was used to isolate animals with targeted events. PCR and genomic sequencing of surrounding regions validated the *Ankle2*^*CRIMIC*^ allele. *Ankle*^*IGFP*^ was generated by RMCE by injecting a plasmid expressing integrase with pBS-KS-attB1-2-PT-SA-SD-EGFP-FlAsH-StrepII-TEV-3xFlag (Nagarkar-Jaiswal *et al.*, 2015b) into *Ankle2*^*CRIMIC*^ animals. Animals with 3XP3-GFP loss were screened using PCR for targeted cassette exchange. Regions flanking the targeted event were sequenced to verify the allele. The *Ankle2-GFPR* was created using recombineering (Venken *et al.*, 2008) of BAC CH321-85N12 (Venken *et al.*, 2006). A GFP donor construct was generated by amplifying the GFP coding region and a selection cassette from plasmid PL-452 C-EGFP with primers containing 50bp homology with the C-terminal end of Ankle2 (5’-GGGATCAACGGTCCTATAACGAGGGGGACACGCCGCTGGGCAATCGGAAC<u>GCAGCCCAATTCCGATCATATTC</u>-3’ and 5’-CATCAATCAGTCGCTGTTTCTGTTTCTGTTTCCGGGCCGATT CCGTTTCA<u>TTACTTGTACAGCTCGTCCATG-3’</u>. Regions matching PL-452 C-EGFP are underlined. CH321-85N12 was transformed into DY380 cells using electroporation. Stable colonies were grown overnight at 30**°**C, induced for recombination functions at 42**°**C for 15 min, and transformed using electroporation (1.8kV, 200Ohm, 25μFD) with the amplified donor construct. Colonies were selected for both the BAC (chloramphenicol) and insertion of GFP (kanamycin). Resulting colonies were verified using PCR, restriction enzyme digestion, and sequencing. The GFP tag selection cassette was removed using Cre mediated excision by transforming the *Ankle2-GFP* BAC into induced EL350 cells. Properly excised events were verified by PCR, absence of growth on kanamycin selection plates, and sequencing.

#### Generation of human ANKLE2 expression constructs

NEB Q5 Site-directed mutagenesis was performed on *P{UASt-hANKLE2}.* Each plasmid was sequence verified and injected into VK37 flies with a plasmid expressing integrase for site-specific integration.

#### Brain immunostaining

Late 3^rd^ instar (based on gut clearance and extruding spiracles) larval brains were dissected in PBS and fixed with 4% PFA/PBS/0.3%Triton for 20 minutes. For immunostaining, brains were blocked in PBS/0.3%Triton/1%BSA/5% normal goat serum and incubated in primary antibody in PBS/0.3%Triton/1%BSA overnight. Primary antibodies include rat anti-Deadpan (Abcam Cat# ab195172, 1:250 or 1:500), mouse anti-Prospero MR1A (Developmental Studies Hybridoma Bank, 1:1000), rat anti-Miranda (1:500, Abcam Cat# ab197788), rabbit anti-aPKC (1:1000, PKCz (C-20) Santa Cruz, discontinued), rabbit anti-GFP (1:1000, Invitrogen Cat# A11122), mouse anti-Calnexin 99a (Developmental Studies Hybridoma Bank, 1:100), mouse anti-Lamin Dm0 ADL67.10 (Developmental Studies Hybridoma Bank, 1:250), guinea pig anti-Bazooka (1:1000) (Siller *et al*., 2006), rat anti-Par6 (1:50) (Rolls *et al.*, 2003), rabbit anti-phospho-Histone H3 (1:1000, Millipore Cat# 06-570), rabbit anti-Ball (1:1000) (Yakulov *et al.*, 2014), rabbit anti-VRK1 (1:1000, Abcam Cat# ab151706), rabbit anti-CNN (1:1000) (Lucas and Raff, 2007), and mouse anti-Strep (Qiagen Cat# 34850, 1:500) with goat or donkey secondary antibodies from Jackson ImmunoResearch used 1:500. Brains were mounted with double sided tape spacers and imaged using a Leica Sp8 with 2 µm or 3 µm sections through the entire brain lobe.

#### Live imaging

3^rd^ instar larvae were dissected in sterile PBS supplemented with 1% FBS and 0.5mM ascorbic acid, fine dissected on an inverted Sarstedt lumox dish 50 in a petroleum jelly well. Samples were imaged on a Leica Sp8 with optimized settings for high quality images without bleaching or a Zeiss 880 with Airy scan (wild type Ankle2-GFP and Sec61β colocalizaiton).

#### Protein immunoprecipitation, mass spectrometry, and western analysis

3^rd^ instar larvae or dissected larval brains from *l(2)glMI07575-GFSTF.0* animals were dissociated in 0.1% CHAPS buffer supplemented with protease and phosphatase inhibitors for at least 30 min on ice, centrifuged for 10 min at 4°C, and supernatant was used for immunoprecipitation or western analysis. For immunopreciptation, 25ul of Allele Biotechnology GFP nanoantibody agarose (nAb, Cat# ABP-NAB-GFPA100) was equilibrated and incubated with lysate 2hrs - overnight at 4°C with rotation. Agarose was spun down for 1 min at 1000 × *g* at 4°C, supernatant was removed, and pellet was washed 3X (1X binding buffer (10mM Tris-HCl pH 7.5,150mM NaCl), 2X wash buffer (10mM Tris-HCl pH7.5, 500mM NaCl)). Remaining agarose pellet was submitted to MD Anderson Proteomics and Metabolomics Facility core for mass spectrometry target identification, or eluted for western analysis in loading buffer. For western analysis, larval brains were dissected and dissociated as stated above, and were lysed in 0.1% CHAPS buffer [[50mM Nacl, 200mM HEPES, 1mM EDTA and protease inhibitor cocktail (Roche)] Loading input was adjusted for brain size and protein concentration. Primary antibodies include rabbit anti-GFP (1:2500, Invitrogen Cat# A11122), rabbit anti-Ball (1:1000) (Herzig *et al.*, 2014), rat anti-L(2)gl (Peng *et al.*, 2000), rabbit anti-aPKC c-20 (1:1000, Santa Cruz, discontinued), rabbit anti-aPKC phosphoT410 (1:1000, Santa Cruz, discontinued), and mouse anti-Actin-c4 (1:5000, Millipore Cat# MAB1501). Secondary antibodies include Rockland DyLight 600 and 800 (1:1000), BioRad Star Bright Blue 700 (1:1000) and Jackson ImmunoResearch HRP conjugated (1:5000). Blots were imaged on a Bio-Rad ChemiDocMP.

#### Human cell immunohistochemistry

Cells were detached using Trypsin-EDTA 0.05% and plated onto 18mm glass cover slips in 6 well plates. Cells were cultured for an additional 3 days under the same conditions before fixing and staining. Cells were rinsed with PBS followed by fixing in 4% paraformaldehyde in PBS. Cells were rinsed and washed 3x in PBST, washed 2x in PBST + 1% BSA (PBSTB), and then blocked in PBSTB + 5% normal goat serum. They were then incubated in PBSTB and primary antibody overnight at 4 degrees Celsius. Cells were then washed 3x in PBSTB, incubated in anti-rabbit Cy5 secondary antibody (1:500) for 2 hours, and washed 3x in PBST. The cells were given a final wash in PBST with DAPI (1:1000) for 30m before mounting using SlowFade glow on glass slides and sealing with nail polish.

#### Exome and Sanger Sequencing

Exome sequencing was performed under the Baylor Hopkins Center for Mendelian Genomics (BHCMG) research initiative as previously described (Lupski *et al.*, 2013). Exome capture was performed with Nimblegen reagents and a custom capture reagent, VCRome2.1. Raw data was processed using the Mercury pipeline, available on DNANexus (http://blog.dnanexus.com/2013-10-22-run-mercury-variant-calling-pipeline/) (Reid *et al.*, 2014) and the ATLAS2 method was used for variant calling followed by an in-house Cassandra annotation pipeline based on Annotation of Genetic Variants (ANNOVAR). The *LLGL1* variant was orthogonally validated and segregated with disease by dideoxy Sanger sequencing of PCR amplicons (Sanger *et al*., 1977).

### QUANTIFICATION AND STATISTICAL ANALYSIS

#### Brain volume

Brains from third instar larvae were stained and mounted with tape spacers and imaged using a Leica Sp8 with 2 µm or 3 µm sections through the entire brain lobe. Resulting stacks were analyzed using the Surfaces function in Imaris (Bitplane) to quantify brain lobe volume as total microns cubed. One lobe from each brain was imaged and a total of 5-10 brains were analyzed per genotype or condition. Brain lobe volumes are displayed as box plots with hinges representing the 25^th^ to 75^th^ percentiles, a line represents the median, and whiskers represent min to max. Statistical significance was determined using one-way ANOVA with multiple comparisons post-test calculated using GraphPad Prism. Brain volumes in Figure 1 are normalized to wild type (*FRT19a*). Average volume from wild type is set to 100%, and each mutant or condition is normalized as percentage of wild type volume. Brain volumes from Figure 4-6 are displayed as total brain volume (μm ^3^).

#### Asymmetric phenotypes

3^rd^ instar larvae were immunostained for pH3, Baz, Par6, aPKC, or Mira as described above. Metaphase neuroblasts (pH3 positive, chromosomes aligned at the metaphase plate) were imaged on a Leica Sp8 (63X). Only metaphase neuroblasts in the correct plane for imaging were analyzed. Mild disruption refers to weak or incomplete crescent localization, and strong disruption indicates no crescent localization. To quantify spindle orientation, CNN was used to mark the plane of division, and aPKC, and Mira were used to establish cortex polarity. Only metaphase neuroblasts in the correct plane for imaging were analyzed. The angle between spindle orientation and cortex polarity was measured using the angle function of ImageJ. Phenotypes are portrayed as percentage of total counted metaphase neuroblasts. For all samples n>20 but <60.

#### VRK1 intensity

Human fibroblasts were stained as described above, imaged on a Zeiss 710 as Z-stacks with equivalent laser power and confocal settings in the same imaging session. Resulting images were analyzed in Imaris (Bitplane) using the Surfaces function to mark nuclear volume. Total intensity sum of the VRK1 channel within the nucleus and nuclear volume were recorded. VRK1 intensity is displayed as intensity sum normalized to volume. One-way ANOVA with multi-comparisons post-test from GraphPad Prism was used to assess significance. Each dot represents one nucleus. Three fields from each cell line were assessed.

## Supplemental Figures

**Figure S1: *ANKLE2* family pedigrees. Related to Figure 1.** Pedigrees and segregation of identified variants for families in which biallelic segregating variants in *ANKLE2* were found. Filled shapes represent affected individuals. Name of the genes, nucleotide changes, protein changes, and respective variants (red) are written below the pedigrees. Filled in arrows designate the proband.

**Figure S2: *Ankle2***^***A***^ **mutations are temperature sensitive. Related to Figure 2.** Wild type (*FRT19a*) and *Ankle2*^*A*^ animals were raised at 18°C, 25°C, or 29°C, and viability and brain volume was assessed. A portion of *Ankle2*^*A*^ mutants raised at 18°C are viable as adults and 3^rd^ instar larval brain volumes are not different from wild type animals. *Ankle2*^*A*^ mutants raised at 25°C or 29°C are lethal and have severely reduce brain volumes.

**Figure S3: Ankle2**^**IGFP**^ **is expressed in the brain. Related to Figure 3.** Single slice from a 3^rd^ instar larval brain from *Ankle2*^*IGFP*^ heterozygous animals stained for GFP (green) and DNA (blue). Ankle2^IGFP^ is expressed throughout the brain.

**Figure S4: Mutations in *ANKLE2* do not alter overall VRK1 protein levels. Related to Figure 5.** Immunostaining of VRK1 and DNA (shown in Figure 5) was used to quantify total VRK1 intensity per cell. Imaris was used to outline cells and intensity sum was assessed and plotted.

**Figure S5: *VRK1* is associated with disease. Related to Figure 5**. Pedigrees and segregation of identified variants for families in which biallelic segregating variants in *VRK1* were found. Filled shapes represent affected individuals. Name of the genes, nucleotide changes, protein changes, and respective variants (red) are written below the pedigrees. Filled in arrows designate the proband. Protein structure with variants is shown below.

**Figure S6: *VRK2* and *VRK3*, paralogs of *VRK1*, are associated with disease. Related to Figure 5.** Pedigrees and segregation of identified variants for families in which biallelic segregating variants in VRK2 and VRK3 were found. Filled shapes represent affected individuals. Name of the genes, nucleotide changes, protein changes, and respective variants (red) are written below the pedigrees. Filled in arrows designate the proband. Sanger traces for families with variants identified in a single gene are shown below the relevant pedigree. Absence of Heterozygosity (AOH) maps for probands is shown below the family of BAB7812. Vertical red line delineates the genes of interest and regions with a bold blue line represent regions of AOH. Protein structures with variants are shown below.

**Figure S7: Variants in the PAR complex and *LLGL1* are associated with disease. Related to Figure 6.** Pedigrees and segregation of identified variants for families in which biallelic segregating variants the PAR complex encoding genes and their paralogs were found. Filled shapes represent affected individuals. Name of the genes, nucleotide changes, protein changes, and respective variants (red) are written below the pedigrees. Filled in arrows designate the proband. Sanger traces for families with variants identified in a single gene are shown below the relevant pedigree.

## Supplemental Tables

**Table S1: Summary of *ANKLE2* and *VRK1* published cases. Related to Figure 1 and 5.**

Published patient variants in ANKLE2 and VRK1 with associated phenotypes.

**Table S2: Summary of *ANKLE2, VRK1, PAR* complex, and *LLGL1* variants in BGL and BHCMG databases. Related to Figure 1, 5, and 6.** Unpublished human variants in the ANKLE2 pathway with analysis of variants. Zyg: Zygosity; vR/tR: Variant Reads to Total Reads Ratio; phyloP: evolutionary conservation score; ARIC: frequency in Atherosclerosis Risk in Communities database; CADD: Combined Annotation Dependent Depletion Score; gnomAD: frequency of heterozygotes and homozygotes in the publicly available gnomAD database; AOH block: Absence of Heterozygosity regions in which homozygous variants were found.

## Supplemental Movies

**Movie S1: Ankle2-GFPR localizes to the nuclear envelope during division. Related to Figure 3**. Time lapse images of live 3^rd^ instar larval brains from *Ankle2-GFPR* animals showing subcellular localization of Ankle2 protein (green). Large cells are neuroblasts. Ankle2 is cytoplasmic during interphase but is recruited to the nuclear envelope during cell division.

**Movie S2: Ankle2-GFPR localizes to the nuclear envelope during division. Related to Figure 3.** Time lapse images of live 3^rd^ instar larval brains from *Ankle2-GFPR* animals showing subcellular localization of Ankle2 protein (green). Large cells are neuroblasts. Ankle2 is cytoplasmic during interphase but is recruited to the nuclear envelope during cell division.

**Movie S3: Miranda localizes to the basal membrane in wild type dividing neuroblasts. Related to Figure 4.** Time lapse imaging of live 3^rd^ instar larval brains from wild type animals expressing Miranda-GFP (green) and Histone-RFP (DNA, white). Large cells are neuroblasts. Note the localization of Miranda at the basal membrane during mitosis.

**Movie S4: *Ankle2***^***A***^ **mutant neuroblasts fail to segregate DNA. Related to Figure 4.** Time lapse imaging of live 3^rd^ instar larval brains from *Ankle2*^*A*^ animals expressing Miranda-GFP (green) and Histone-RFP (DNA, white). Large cells are neuroblasts. Note reduced localization of Miranda at the basal membrane and DNA segregation failures during mitosis.

**Movie S5: *Ankle2***^***A***^ **mutant neuroblasts fail to undergo cytokinesis. Related to Figure 4.** Time lapse imaging of live 3^rd^ instar larval brains from *Ankle2*^*A*^ animals expressing Miranda-GFP (green). Large cells are neuroblasts. Note reduced localization of Miranda at the basal membrane and cytokinesis failure during mitosis.

**Movie S6: *Ankle2***^***A***^ **mutant neuroblasts have defective Miranda localization. Related to Figure 4.** Time lapse imaging of live 3^rd^ instar larval brains from *Ankle2*^*A*^ animals expressing Miranda-GFP (green) and Histone-RFP (DNA, white). Large cells are neuroblasts. Note reduced localization of Miranda at the basal membrane.

**Movie S7: Ballchen localizes to the nucleus during interphase and redistributes throughout the entire cell during mitosis. Related to Figure 5.** Time lapse imaging of live 3^rd^ instar larval brains from wild type animals expressing Ballchen-GFP (green). Large cells are neuroblasts. Note the nuclear localization of Ball during interphase, redistribution during mitosis, and recruitment back to the nucleus upon completion of mitosis.

## REFERENCES

Andersen, R. O., Turnbull, D. W., Johnson, E. A. and Doe, C. Q. (2012) ‘Sgt1 acts via an LKB1/AMPK pathway to establish cortical polarity in larval neuroblasts’, Dev Biol, 363(1), pp. 258–65.

Asencio, C., Davidson, I. F., Santarella-Mellwig, R., Ly-Hartig, T. B., Mall, M., Wallenfang, M. R., Mattaj, I. W. and Gorjanacz, M. (2012) ‘Coordination of kinase and phosphatase activities by Lem4 enables nuclear envelope reassembly during mitosis’, Cell, 150(1), pp. 122–35.

Atwood, S. X., Chabu, C., Penkert, R. R., Doe, C. Q. and Prehoda, K. E. (2007) ‘Cdc42 acts downstream of Bazooka to regulate neuroblast polarity through Par-6 aPKC’, J Cell Sci, 120(Pt 18), pp. 3200–6.

Atwood, S. X. and Prehoda, K. E. (2009) ‘aPKC phosphorylates Miranda to polarize fate determinants during neuroblast asymmetric cell division’, Curr Biol, 19(9), pp. 723–9.

Ayoub, A. E., Oh, S., Xie, Y., Leng, J., Cotney, J., Dominguez, M. H., Noonan, J. P. and Rakic, P. (2011) ‘Transcriptional programs in transient embryonic zones of the cerebral cortex defined by high-resolution mRNA sequencing’, Proc Natl Acad Sci U S A, 108(36), pp. 14950–5.

Bamshad, M. J., Shendure, J. A., Valle, D., Hamosh, A., Lupski, J. R., Gibbs, R. A., Boerwinkle, E., Lifton, R. P., Gerstein, M., Gunel, M., Mane, S., Nickerson, D. A. and Genomics, C. f. M. (2012) ‘The Centers for Mendelian Genomics: a new large-scale initiative to identify the genes underlying rare Mendelian conditions’, Am J Med Genet A, 158A(7), pp. 1523–5.

Barton, L. J., Soshnev, A. A. and Geyer, P. K. (2015) ‘Networking in the nucleus: a spotlight on LEM-domain proteins’, Curr Opin Cell Biol, 34, pp. 1–8.

Betschinger, J., Mechtler, K. and Knoblich, J. A. (2003) ‘The Par complex directs asymmetric cell division by phosphorylating the cytoskeletal protein Lgl’, Nature, 422(6929), pp. 326–30.

Bier, E., Vaessin, H., Younger-Shepherd, S., Jan, L. Y. and Jan, Y. N. (1992) ‘deadpan, an essential pan-neural gene in Drosophila, encodes a helix-loop-helix protein similar to the hairy gene product’, Genes Dev, 6(11), pp. 2137–51.

Bischof, J., Björklund, M., Furger, E., Schertel, C., Taipale, J. and Basler, K. (2013) ‘A versatile platform for creating a comprehensive UAS-ORFeome library in Drosophila’, Development, 140(11), pp. 2434–42.

Bonaccorsi, S., Mottier, V., Giansanti, M. G., Bolkan, B. J., Williams, B., Goldberg, M. L. and Gatti, M. (2007) ‘The Drosophila Lkb1 kinase is required for spindle formation and asymmetric neuroblast division’, Development, 134(11), pp. 2183–93.

Brunetti-Pierri, N., Berg, J. S., Scaglia, F., Belmont, J., Bacino, C. A., Sahoo, T., Lalani, S. R., Graham, B., Lee, B., Shinawi, M., Shen, J., Kang, S. H., Pursley, A., Lotze, T., Kennedy, G., Lansky-Shafer, S., Weaver, C., Roeder, E. R., Grebe, T. A., Arnold, G. L., Hutchison, T., Reimschisel, T., Amato, S., Geragthy, M. T., Innis, J. W., Obersztyn, E., Nowakowska, B., Rosengren, S. S., Bader, P. I., Grange, D. K., Naqvi, S., Garnica, A. D., Bernes, S. M., Fong, C. T., Summers, A., Walters, W. D., Lupski, J. R., Stankiewicz, P., Cheung, S. W. and Patel, A. (2008) ‘Recurrent reciprocal 1q21.1 deletions and duplications associated with microcephaly or macrocephaly and developmental and behavioral abnormalities’, Nat Genet, 40(12), pp. 1466–71.

Cabernard, C. and Doe, C. Q. (2009) ‘Apical/basal spindle orientation is required for neuroblast homeostasis and neuronal differentiation in Drosophila’, Dev Cell, 17(1), pp. 134–41.

Campbell, G., Göring, H., Lin, T., Spana, E., Andersson, S., Doe, C. Q. and Tomlinson, A. (1994) ‘RK2, a glial-specific homeodomain protein required for embryonic nerve cord condensation and viability in Drosophila’, Development, 120(10), pp. 2957–66.

Carvalho, C. A., Moreira, S., Ventura, G., Sunkel, C. E. and Morais-de-Sá, E. (2015) ‘Aurora A triggers Lgl cortical release during symmetric division to control planar spindle orientation’, Curr Biol, 25(1), pp. 53–60.

Chabu, C. and Doe, C. Q. (2008) ‘Dap160/intersectin binds and activates aPKC to regulate cell polarity and cell cycle progression’, Development, 135(16), pp. 2739–46.

Chabu, C. and Doe, C. Q. (2009) ‘Twins/PP2A regulates aPKC to control neuroblast cell polarity and self-renewal’, Dev Biol, 330(2), pp. 399–405.

Clarkson, M. and Saint, R. (1999) ‘A His2AvDGFP fusion gene complements a lethal His2AvD mutant allele and provides an in vivo marker for Drosophila chromosome behavior’, DNA Cell Biol, 18(6), pp. 457–62.

Cullen, C. F., Brittle, A. L., Ito, T. and Ohkura, H. (2005) ‘The conserved kinase NHK-1 is essential for mitotic progression and unifying acentrosomal meiotic spindles in Drosophila melanogaster’, J Cell Biol, 171(4), pp. 593–602.

Devhare, P., Meyer, K., Steele, R., Ray, R. B. and Ray, R. (2017) ‘Zika virus infection dysregulates human neural stem cell growth and inhibits differentiation into neuroprogenitor cells’, Cell Death Dis, 8(10), pp. e3106.

Drier, E. A., Tello, M. K., Cowan, M., Wu, P., Blace, N., Sacktor, T. C. and Yin, J. C. (2002) ‘Memory enhancement and formation by atypical PKM activity in Drosophila melanogaster’, Nat Neurosci, 5(4), pp. 316–24.

Dumas, L. J., O’Bleness, M. S., Davis, J. M., Dickens, C. M., Anderson, N., Keeney, J. G., Jackson, J., Sikela, M., Raznahan, A., Giedd, J., Rapoport, J., Nagamani, S. S., Erez, A., Brunetti-Pierri, N., Sugalski, R., Lupski, J. R., Fingerlin, T., Cheung, S. W. and Sikela, J. M. (2012) ‘DUF1220-domain copy number implicated in human brain-size pathology and evolution’, Am J Hum Genet, 91(3), pp. 444–54.

Ferrari, S., Marin, O., Pagano, M. A., Meggio, F., Hess, D., El-Shemerly, M., Krystyniak, A. and Pinna, L. A. (2005) ‘Aurora-A site specificity: a study with synthetic peptide substrates’, Biochem J, 390(Pt 1), pp. 293–302.

Gallaud, E., Pham, T. and Cabernard, C. (2017) ‘Drosophila melanogaster Neuroblasts: A Model for Asymmetric Stem Cell Divisions’, Results Probl Cell Differ, 61, pp. 183–210.

Gateff, E. and Schneiderman, H. A. (1974) ‘Developmental capacities of benign and malignant neoplasms ofDrosophila’, Wilhelm Roux Arch Entwickl Mech Org, 176(1), pp. 23–65.

Gonzaga-Jauregui, C., Lotze, T., Jamal, L., Penney, S., Campbell, I. M., Pehlivan, D., Hunter, J. V., Woodbury, S. L., Raymond, G., Adesina, A. M., Jhangiani, S. N., Reid, J. G., Muzny, D. M., Boerwinkle, E., Lupski, J. R., Gibbs, R. A. and Wiszniewski, W. (2013) ‘Mutations in VRK1 associated with complex motor and sensory axonal neuropathy plus microcephaly’, JAMA Neurol, 70(12), pp. 1491–8.

Greenberg, F., Guzzetta, V., Montes de Oca-Luna, R., Magenis, R. E., Smith, A. C., Richter, S. F., Kondo, I., Dobyns, W. B., Patel, P. I. and Lupski, J. R. (1991) ‘Molecular analysis of the Smith-Magenis syndrome: a possible contiguous-gene syndrome associated with del(17)(p11.2)’, Am J Hum Genet, 49(6), pp. 1207–18.

Greenberg, F., Lewis, R. A., Potocki, L., Glaze, D., Parke, J., Killian, J., Murphy, M. A., Williamson, D., Brown, F., Dutton, R., McCluggage, C., Friedman, E., Sulek, M. and Lupski, J. R. (1996) ‘Multi-disciplinary clinical study of Smith-Magenis syndrome (deletion 17p11.2)’, Am J Med Genet, 62(3), pp. 247–54.

Harsh, S., Ozakman, Y., Kitchen, S. M., Paquin-Proulx, D., Nixon, D. F. and Eleftherianos, I. (2018) ‘Dicer-2 Regulates Resistance and Maintains Homeostasis against Zika Virus Infection in’, J Immunol, 201(10), pp. 3058–3072.

Herzig, B., Yakulov, T. A., Klinge, K., Günesdogan, U., Jäckle, H. and Herzig, A. (2014) ‘Bällchen is required for self-renewal of germline stem cells in Drosophila melanogaster’, Biol Open, 3(6), pp. 510–21.

Homem, C. C. and Knoblich, J. A. (2012) ‘Drosophila neuroblasts: a model for stem cell biology’, Development, 139(23), pp. 4297–310.

Ito, K., Awano, W., Suzuki, K., Hiromi, Y. and Yamamoto, D. (1997) ‘The Drosophila mushroom body is a quadruple structure of clonal units each of which contains a virtually identical set of neurones and glial cells’, Development, 124(4), pp. 761–71.

Jayaraman, D., Bae, B. I. and Walsh, C. A. (2018) ‘The Genetics of Primary Microcephaly’, Annu Rev Genomics Hum Genet, 19, pp. 177–200.

Juyal, R. C., Figuera, L. E., Hauge, X., Elsea, S. H., Lupski, J. R., Greenberg, F., Baldini, A. and Patel, P. I. (1996) ‘Molecular analyses of 17p11.2 deletions in 62 Smith-Magenis syndrome patients’, Am J Hum Genet, 58(5), pp. 998–1007.

Karaca, E., Posey, J. E., Coban Akdemir, Z., Pehlivan, D., Harel, T., Jhangiani, S. N., Bayram, Y., Song, X., Bahrambeigi, V., Yuregir, O. O., Bozdogan, S., Yesil, G., Isikay, S., Muzny, D., Gibbs, R. A. and Lupski, J. R. (2018) ‘Phenotypic expansion illuminates multilocus pathogenic variation’, Genet Med, 20(12), pp. 1528–1537.

Kim, S., Gailite, I., Moussian, B., Luschnig, S., Goette, M., Fricke, K., Honemann-Capito, M., Grubmüller, H. and Wodarz, A. (2009) ‘Kinase-activity-independent functions of atypical protein kinase C in Drosophila’, J Cell Sci, 122(Pt 20), pp. 3759–71.

Kondo, S. and Ueda, R. (2013) ‘Highly improved gene targeting by germline-specific Cas9 expression in Drosophila’, Genetics, 195(3), pp. 715–21.

Krahn, M. P., Egger-Adam, D. and Wodarz, A. (2009) ‘PP2A antagonizes phosphorylation of Bazooka by PAR-1 to control apical-basal polarity in dividing embryonic neuroblasts’, Dev Cell, 16(6), pp. 901–8.

Lee, C. Y., Andersen, R. O., Cabernard, C., Manning, L., Tran, K. D., Lanskey, M. J., Bashirullah, A. and Doe, C. Q. (2006) ‘Drosophila Aurora-A kinase inhibits neuroblast self-renewal by regulating aPKC/Numb cortical polarity and spindle orientation’, Genes Dev, 20(24), pp. 3464–74.

Lee, P. T., Zirin, J., Kanca, O., Lin, W. W., Schulze, K. L., Li-Kroeger, D., Tao, R., Devereaux, C., Hu, Y., Chung, V., Fang, Y., He, Y., Pan, H., Ge, M., Zuo, Z., Housden, B. E., Mohr, S. E., Yamamoto, S., Levis, R. W., Spradling, A. C., Perrimon, N. and Bellen, H. J. (2018) ‘A gene-specific *T2A-GAL4* library for *Drosophila*’, Elife, 7.

Lee, T. and Luo, L. (2001) ‘Mosaic analysis with a repressible cell marker (MARCM) for Drosophila neural development’, Trends Neurosci, 24(5), pp. 251–4.

Leitges, M., Sanz, L., Martin, P., Duran, A., Braun, U., García, J. F., Camacho, F., Diaz-Meco, M. T., Rennert, P. D. and Moscat, J. (2001) ‘Targeted disruption of the zetaPKC gene results in the impairment of the NF-kappaB pathway’, Mol Cell, 8(4), pp. 771–80.

Li, C., Xu, D., Ye, Q., Hong, S., Jiang, Y., Liu, X., Zhang, N., Shi, L., Qin, C. F. and Xu, Z. (2016) ‘Zika Virus Disrupts Neural Progenitor Development and Leads to Microcephaly in Mice’, Cell Stem Cell, 19(5), pp. 672.

Liburd, N., Ghosh, M., Riazuddin, S., Naz, S., Khan, S., Ahmed, Z., Liang, Y., Menon, P. S., Smith, T., Smith, A. C., Chen, K. S., Lupski, J. R., Wilcox, E. R., Potocki, L. and Friedman, T. B. (2001) ‘Novel mutations of MYO15A associated with profound deafness in consanguineous families and moderately severe hearing loss in a patient with Smith-Magenis syndrome’, Hum Genet, 109(5), pp. 535–41.

Lin, F., Blake, D. L., Callebaut, I., Skerjanc, I. S., Holmer, L., McBurney, M. W., Paulin-Levasseur, M. and Worman, H. J. (2000) ‘MAN1, an inner nuclear membrane protein that shares the LEM domain with lamina-associated polypeptide 2 and emerin’, J Biol Chem, 275(7), pp. 4840–7.

Lin, M. Y., Wang, Y. L., Wu, W. L., Wolseley, V., Tsai, M. T., Radic, V., Thornton, M. E., Grubbs, B. H., Chow, R. H. and Huang, I. C. (2017) ‘Zika Virus Infects Intermediate Progenitor Cells and Post-mitotic Committed Neurons in Human Fetal Brain Tissues’, Sci Rep, 7(1), pp. 14883.

Liu, Y., Gordesky-Gold, B., Leney-Greene, M., Weinbren, N. L., Tudor, M. and Cherry, S. (2018) ‘Inflammation-Induced, STING-Dependent Autophagy Restricts Zika Virus Infection in the Drosophila Brain’, Cell Host Microbe, 24(1), pp. 57–68.e3.

Lucas, E. P. and Raff, J. W. (2007) ‘Maintaining the proper connection between the centrioles and the pericentriolar matrix requires Drosophila centrosomin’, J Cell Biol, 178(5), pp. 725–32.

Luo, L., Liao, Y. J., Jan, L. Y. and Jan, Y. N. (1994) ‘Distinct morphogenetic functions of similar small GTPases: Drosophila Drac1 is involved in axonal outgrowth and myoblast fusion’, Genes Dev, 8(15), pp. 1787–802.

Lupski, J. R. (2015) ‘Structural variation mutagenesis of the human genome: Impact on disease and evolution’, Environ Mol Mutagen, 56(5), pp. 419–36.

Lupski, J. R., Gonzaga-Jauregui, C., Yang, Y., Bainbridge, M. N., Jhangiani, S., Buhay, C. J., Kovar, C. L., Wang, M., Hawes, A. C., Reid, J. G., Eng, C., Muzny, D. M. and Gibbs, R. A. (2013) ‘Exome sequencing resolves apparent incidental findings and reveals further complexity of SH3TC2 variant alleles causing Charcot-Marie-Tooth neuropathy’, Genome Med, 5(6), pp. 57.

Manfruelli, P., Arquier, N., Hanratty, W. P. and Sémériva, M. (1996) ‘The tumor suppressor gene, lethal(2)giant larvae (1(2)g1), is required for cell shape change of epithelial cells during Drosophila development’, Development, 122(7), pp. 2283–94.

Marchler-Bauer, A., Bo, Y., Han, L., He, J., Lanczycki, C. J., Lu, S., Chitsaz, F., Derbyshire, M. K., Geer, R. C., Gonzales, N. R., Gwadz, M., Hurwitz, D. I., Lu, F., Marchler, G. H., Song, J. S., Thanki, N., Wang, Z., Yamashita, R. A., Zhang, D., Zheng, C., Geer, L. Y. and Bryant, S. H. (2017) ‘CDD/SPARCLE: functional classification of proteins via subfamily domain architectures’, Nucleic Acids Res, 45(D1), pp. D200–D203.

Mlakar, J., Korva, M., Tul, N., Popović, M., Poljšak-Prijatelj, M., Mraz, J., Kolenc, M., Resman Rus, K., Vesnaver Vipotnik, T., Fabjan Vodušek, V., Vizjak, A., Pižem, J., Petrovec, M. and Avšič Županc, T. (2016) ‘Zika Virus Associated with Microcephaly’, N Engl J Med, 374(10), pp. 951–8.

Moore, C. A., Staples, J. E., Dobyns, W. B., Pessoa, A., Ventura, C. V., Fonseca, E. B., Ribeiro, E. M., Ventura, L. O., Neto, N. N., Arena, J. F. and Rasmussen, S. A. (2017) ‘Characterizing the Pattern of Anomalies in Congenital Zika Syndrome for Pediatric Clinicians’, JAMA Pediatr, 171(3), pp. 288–295.

Nagarkar-Jaiswal, S., DeLuca, S. Z., Lee, P. T., Lin, W. W., Pan, H., Zuo, Z., Lv, J., Spradling, A. C. and Bellen, H. J. (2015a) ‘A genetic toolkit for tagging intronic MiMIC containing genes’, Elife, 4.

Nagarkar-Jaiswal, S., Lee, P. T., Campbell, M. E., Chen, K., Anguiano-Zarate, S., Gutierrez, M. C., Busby, T., Lin, W. W., He, Y., Schulze, K. L., Booth, B. W., Evans-Holm, M., Venken, K. J., Levis, R. W., Spradling, A. C., Hoskins, R. A. and Bellen, H. J. (2015b) ‘A library of MiMICs allows tagging of genes and reversible, spatial and temporal knockdown of proteins in Drosophila’, Elife, 4.

Nam, S. C., Mukhopadhyay, B. and Choi, K. W. (2007) ‘Antagonistic functions of Par-1 kinase and protein phosphatase 2A are required for localization of Bazooka and photoreceptor morphogenesis in Drosophila’, Dev Biol, 306(2), pp. 624–35.

Nichols, R. J. and Traktman, P. (2004) ‘Characterization of three paralogous members of the Mammalian vaccinia related kinase family’, J Biol Chem, 279(9), pp. 7934–46.

Peng, C. Y., Manning, L., Albertson, R. and Doe, C. Q. (2000) ‘The tumour-suppressor genes lgl and dlg regulate basal protein targeting in Drosophila neuroblasts’, Nature, 408(6812), pp. 596–600.

Petronczki, M. and Knoblich, J. A. (2001) ‘DmPAR-6 directs epithelial polarity and asymmetric cell division of neuroblasts in Drosophila’, Nat Cell Biol, 3(1), pp. 43–9.

Port, F., Chen, H. M., Lee, T. and Bullock, S. L. (2014) ‘Optimized CRISPR/Cas tools for efficient germline and somatic genome engineering in Drosophila’, Proc Natl Acad Sci U S A.

Posey, J. E., O’Donnell-Luria, A. H., Chong, J. X., Harel, T., Jhangiani, S. N., Coban Akdemir, Z. H., Buyske, S., Pehlivan, D., Carvalho, C. M. B., Baxter, S., Sobreira, N., Liu, P., Wu, N., Rosenfeld, J. A., Kumar, S., Avramopoulos, D., White, J. J., Doheny, K. F., Witmer, P. D., Boehm, C., Sutton, V. R., Muzny, D. M., Boerwinkle, E., Günel, M., Nickerson, D. A., Mane, S., MacArthur, D. G., Gibbs, R. A., Hamosh, A., Lifton, R. P., Matise, T. C., Rehm, H. L., Gerstein, M., Bamshad, M. J., Valle, D., Lupski, J. R. and Genomics, C. f. M. (2019) ‘Insights into genetics, human biology and disease gleaned from family based genomic studies’, Genet Med.

Reid, J. G., Carroll, A., Veeraraghavan, N., Dahdouli, M., Sundquist, A., English, A., Bainbridge, M., White, S., Salerno, W., Buhay, C., Yu, F., Muzny, D., Daly, R., Duyk, G., Gibbs, R. A. and Boerwinkle, E. (2014) ‘Launching genomics into the cloud: deployment of Mercury, a next generation sequence analysis pipeline’, BMC Bioinformatics, 15, pp. 30.

Renbaum, P., Kellerman, E., Jaron, R., Geiger, D., Segel, R., Lee, M., King, M. C. and Levy-Lahad, E. (2009) ‘Spinal muscular atrophy with pontocerebellar hypoplasia is caused by a mutation in the VRK1 gene’, Am J Hum Genet, 85(2), pp. 281–9.

Riedel, F., Gillingham, A. K., Rosa-Ferreira, C., Galindo, A. and Munro, S. (2016) ‘An antibody toolkit for the study of membrane traffic in Drosophila melanogaster’, Biol Open, 5(7), pp. 987–92.

Riemer, D., Stuurman, N., Berrios, M., Hunter, C., Fisher, P. A. and Weber, K. (1995) ‘Expression of Drosophila lamin C is developmentally regulated: analogies with vertebrate A-type lamins’, J Cell Sci, 108 (Pt 10), pp. 3189–98.

Rolls, M. M., Albertson, R., Shih, H. P., Lee, C. Y. and Doe, C. Q. (2003) ‘Drosophila aPKC regulates cell polarity and cell proliferation in neuroblasts and epithelia’, J Cell Biol, 163(5), pp. 1089–98.

Sanger, F., Nicklen, S. and Coulson, A. R. (1977) ‘DNA sequencing with chain-terminating inhibitors’, Proc Natl Acad Sci U S A, 74(12), pp. 5463–7.

Sanz-García, M., Vázquez-Cedeira, M., Kellerman, E., Renbaum, P., Levy-Lahad, E. and Lazo, P. A. (2011) ‘Substrate profiling of human vaccinia-related kinases identifies coilin, a Cajal body nuclear protein, as a phosphorylation target with neurological implications’, J Proteomics, 75(2), pp. 548–60.

Sarov, M., Barz, C., Jambor, H., Hein, M. Y., Schmied, C., Suchold, D., Stender, B., Janosch, S. K J V. V., Krishnan, R. T., Krishnamoorthy, A., Ferreira, I. R., Ejsmont, R. K., Finkl, K., Hasse, S., Kämpfer, P., Plewka, N., Vinis, E., Schloissnig, S., Knust, E., Hartenstein, V., Mann, M., Ramaswami, M., VijayRaghavan, K., Tomancak, P. and Schnorrer, F. (2016) ‘A genome-wide resource for the analysis of protein localisation in Drosophila’, Elife, 5, pp. e12068.

Schober, M., Schaefer, M. and Knoblich, J. A. (1999) ‘Bazooka recruits Inscuteable to orient asymmetric cell divisions in Drosophila neuroblasts’, Nature, 402(6761), pp. 548–51.

Segura-Totten, M., Kowalski, A. K., Craigie, R. and Wilson, K. L. (2002) ‘Barrier-to-autointegration factor: major roles in chromatin decondensation and nuclear assembly’, J Cell Biol, 158(3), pp. 475–85.

Shah, P. S., Link, N., Jang, G. M., Sharp, P. P., Zhu, T., Swaney, D. L., Johnson, J. R., Von Dollen, J., Ramage, H. R., Satkamp, L., Newton, B., Hüttenhain, R., Petit, M. J., Baum, T., Everitt, A., Laufman, O., Tassetto, M., Shales, M., Stevenson, E., Iglesias, G. N., Shokat, L., Tripathi, S., Balasubramaniam, V., Webb, L. G., Aguirre, S., Willsey, A. J., Garcia-Sastre, A., Pollard, K. S., Cherry, S., Gamarnik, A. V., Marazzi, I., Taunton, J., Fernandez-Sesma, A., Bellen, H. J., Andino, R. and Krogan, N. J. (2018) ‘Comparative Flavivirus-Host Protein Interaction Mapping Reveals Mechanisms of Dengue and Zika Virus Pathogenesis’, Cell, 175(7), pp. 1931–1945.e18.

Shaheen, R., Maddirevula, S., Ewida, N., Alsahli, S., Abdel-Salam, G. M. H., Zaki, M. S., Tala, A., Alhashem, A., Softah, A., Al-Owain, M., Alazami, A. M., Abadel, B., Patel, N., Al-Sheddi, Alomar R., Alobeid, E., Ibrahim, N., Hashem, M., Abdulwahab, F., Hamad, M., Tabarki, B., Alwadei, A. H., Alhazzani, F., Bashiri, F. A., Kentab, A., Şahintürk, S., Sherr, E., Fregeau, B., Sogati, S., Alshahwan, S. A. M., Alkhalifi, S., Alhumaidi, Z., Temtamy, S., Aglan, M., Otaify, G., Girisha, K. M., Tulbah, M., Seidahmed, M. Z., Salih, M. A., Abouelhoda, M., Momin, A. A., Saffar, M. A., Partlow, J. N., Arold, S. T., Faqeih, E., Walsh, C. and Alkuraya, F. S. (2018) ‘Genomic and phenotypic delineation of congenital microcephaly’, Genet Med.

Shinawi, M., Liu, P., Kang, S. H., Shen, J., Belmont, J. W., Scott, D. A., Probst, F. J., Craigen, W. J., Graham, B. H., Pursley, A., Clark, G., Lee, J., Proud, M., Stocco, A., Rodriguez, D. L., Kozel, B. A., Sparagana, S., Roeder, E. R., McGrew, S. G., Kurczynski, T. W., Allison, L. J., Amato, S., Savage, S., Patel, A., Stankiewicz, P., Beaudet, A. L., Cheung, S. W. and Lupski, J. R. (2010) ‘Recurrent reciprocal 16p11.2 rearrangements associated with global developmental delay, behavioural problems, dysmorphism, epilepsy, and abnormal head size’, J Med Genet, 47(5), pp. 332–41.

Siller, K. H., Cabernard, C. and Doe, C. Q. (2006) ‘The NuMA-related Mud protein binds Pins and regulates spindle orientation in Drosophila neuroblasts’, Nat Cell Biol, 8(6), pp. 594–600.

Smith, A. C., McGavran, L., Robinson, J., Waldstein, G., Macfarlane, J., Zonona, J., Reiss, J., Lahr, M., Allen, L. and Magenis, E. (1986) ‘Interstitial deletion of (17)(p11.2p11.2) in nine patients’, Am J Med Genet, 24(3), pp. 393–414.

Snyers, L., Erhart, R., Laffer, S., Pusch, O., Weipoltshammer, K. and Schöfer, C. 2018. LEM4/ANKLE-2 deficiency impairs post-mitotic re-localization of BAF, LAP2α and LaminA to the nucleus, causes nuclear envelope instability in telophase and leads to hyperploidy in HeLa cells European Journal of Cell Biology: Elsevier.

Summerville, J. B., Faust, J. F., Fan, E., Pendin, D., Daga, A., Formella, J., Stern, M. and McNew, J. A. (2016) ‘The effects of ER morphology on synaptic structure and function in Drosophila melanogaster’, J Cell Sci, 129(8), pp. 1635–48.

Tang, H., Hammack, C., Ogden, S. C., Wen, Z., Qian, X., Li, Y., Yao, B., Shin, J., Zhang, F., Lee, E. M., Christian, K. M., Didier, R. A., Jin, P., Song, H. and Ming, G. L. (2016) ‘Zika Virus Infects Human Cortical Neural Progenitors and Attenuates Their Growth’, Cell Stem Cell, 18(5), pp. 587–90.

Venken, K. J., He, Y., Hoskins, R. A. and Bellen, H. J. (2006) ‘P[acman]: a BAC transgenic platform for targeted insertion of large DNA fragments in D. melanogaster’, Science, 314(5806), pp. 1747–51.

Venken, K. J., Kasprowicz, J., Kuenen, S., Yan, J., Hassan, B. A. and Verstreken, P. (2008) ‘Recombineering-mediated tagging of Drosophila genomic constructs for in vivo localization and acute protein inactivation’, Nucleic Acids Res, 36(18), pp. e114.

Venken, K. J., Schulze, K. L., Haelterman, N. A., Pan, H., He, Y., Evans-Holm, M., Carlson, J. W., Levis, R. W., Spradling, A. C., Hoskins, R. A. and Bellen, H. J. (2011) ‘MiMIC: a highly versatile transposon insertion resource for engineering Drosophila melanogaster genes’, Nat Methods, 8(9), pp. 737–43.

Wang, C., Chang, K. C., Somers, G., Virshup, D., Ang, B. T., Tang, C., Yu, F. and Wang, H. (2009) ‘Protein phosphatase 2A regulates self-renewal of Drosophila neural stem cells’, Development, 136(13), pp. 2287–96.

Wang, H., Ouyang, Y., Somers, W. G., Chia, W. and Lu, B. (2007) ‘Polo inhibits progenitor self-renewal and regulates Numb asymmetry by phosphorylating Pon’, Nature, 449(7158), pp. 96–100.

Wang, H., Somers, G. W., Bashirullah, A., Heberlein, U., Yu, F. and Chia, W. (2006) ‘Aurora-A acts as a tumor suppressor and regulates self-renewal of Drosophila neuroblasts’, Genes Dev, 20(24), pp. 3453–63.

Wirtz-Peitz, F., Nishimura, T. and Knoblich, J. A. (2008) ‘Linking cell cycle to asymmetric division: Aurora-A phosphorylates the Par complex to regulate Numb localization’, Cell, 135(1), pp. 161–73.

Wodarz, A., Hinz, U., Engelbert, M. and Knust, E. (1995) ‘Expression of crumbs confers apical character on plasma membrane domains of ectodermal epithelia of Drosophila’, Cell, 82(1), pp. 67–76.

Yakulov, T., Günesdogan, U., Jäckle, H. and Herzig, A. (2014) ‘Bällchen participates in proliferation control and prevents the differentiation of Drosophila melanogaster neuronal stem cells’, Biol Open, 3(10), pp. 881–6.

Yamamoto, S., Jaiswal, M., Charng, W. L., Gambin, T., Karaca, E., Mirzaa, G., Wiszniewski, W., Sandoval, H., Haelterman, N. A., Xiong, B., Zhang, K., Bayat, V., David, G., Li, T., Chen, K., Gala, U., Harel, T., Pehlivan, D., Penney, S., Vissers, L. E., de Ligt, J., Jhangiani, S. N., Xie, Y., Tsang, S. H., Parman, Y., Sivaci, M., Battaloglu, E., Muzny, D., Wan, Y. W., Liu, Z., Lin-Moore, A. T., Clark, R. D., Curry, C. J., Link, N., Schulze, K. L., Boerwinkle, E., Dobyns, W. B., Allikmets, R., Gibbs, R. A., Chen, R., Lupski, J. R., Wangler, M. F. and Bellen, H. J. (2014) ‘A drosophila genetic resource of mutants to study mechanisms underlying human genetic diseases’, Cell, 159(1), pp. 200–14.

