## Supplemental Information for "Ankle2, A Target of Zika Virus, Controls Asymmetric Cell Division of Neuroblasts and Uncovers a Novel Microcephaly Pathway"

<u>Supplemental Clinical Information</u>

**BAB4821**

A homozygous missense variant in *VRK2* (c.1234G>A;p.D412N; **Table S2, Figure S6**) was observed in a proband, BAB4821 born to consanguineous Turkish parents, with nanophthalmos. The variant was in a region of absence of heterozygosity (AOH) ~7.7Mb in length gleaned from unphased ES data (Karaca *et al.*, 2018) and consistent with identity-by-descent.

**BAB7812**

In BAB7812, a patient with severe microcephaly (Z = -6.7) and additional brain malformations including cortical dysplasia and agenesis of the corpus callosum, a homozygous variant with a predicted detrimental effect on splicing in a canonical splice site within *VRK3* (c.139+2 T>G) was found (**Table S2**). Additional homozygous variants were found in the same (*PNKP*) and an adjacent (*RNT2*) AOH block as *VRK3* (**Table S2**). All three homozygous variants are predicted deleterious. *PNKP* has a known disease association with microcephaly, seizures, and developmental delay (MIM# 613402) while *RNT2* has a known disease association with Spastic paraplegia 12 (MIM# 604805). Multilocus variation may contribute to the phenotype in BAB7812. The proband was born full term to consanguineous Turkish parents. At 5 months of age, he developed seizures. At 32 months, he was unable to walk, and his weight was 9.8kg (-3.34SD), height was 85cm (-2.03SD), and head circumference was 36.7cm (-6.7SD). Physical examination revealed thick eyebrows, protuberant ears, and almond shaped eyes. Metabolic and ocular examinations were within normal limits. He received continued follow-up care for microcephaly and refractory epilepsy, and special education for associated intellectual disability and developmental delay. His occipital frontal circumference was below the

3rd percentile when measured at 8-11 years of age. Parents and an unaffected sibling were heterozygous for the variant, demonstrating segregation of the variant with disease.

**BAB10531**

A BHCMG search database found a homozygous predicted deleterious missense variant in *PARD3B* (c.1222G>A, p.G408S) in BAB10531 who has microcephaly (**Table S2, Figure S7**). At 15 months, weight he was 9kg (-2.1SD), height was 79cm (0.4SD), and head circumference was 44.5cm (Z = -2.2 measured at 15m and and 18m). Both unaffected consanguineous parents of Turkish descent were heterozygous for the variant that mapped to a region of AOH ~22Mb in length. BAB10531 has had frequent upper respiratory tract infections and was admitted at 17 months of age to investigate a cause for with no clear etiology found. Follow-up head circumference was performed at 18 months, which demonstrated microcephaly was non-progressive (45cm, -2.2SD). Notably, no follow-up for intellectual disability was described on recruitment to the BHCMG.

**BAB8223**

In BAB8223, a proband with distal arthrogryposis, a homozygous missense variant in *PARD3B* (c.1654G>A;p.A552T) which segregated with disease was found (**Table S2, Figure S7)**. The proband was born to consanguineous Turkish parents and the homozygous variant was found within a region of AOH ~37Mb in length. At age 5.5y head circumference was 49cm (13th percentile, ~ -1SD). Cognitive and motor development is within normal limits.

**BAB701**

BAB701 has phenotype consistent with SMS, including brachycephaly, broad face, frontal bossing, downturned mouth, sleep disturbance, mild ventriculomegaly, and intellectual disability. The proband carries a deletion of 17p11.2 suggested to be of maternal origin (Juyal *et* *al.*, 1996). Notably the common deletion region in SMS (17p11.2) includes *LLGL1*. In addition to the common deletion of 17p11.2, the proband has a hemizygous predicted deleterious missense variant in *LLGL1* (c.725G>A, pS242N) (**Table S2, Figure S7**). While most of the proband's phenotypic features are consistent with an SMS phenotype, he was also found to have colobomatous microphthalmia. This trait is not typical of SMS, and while up to 85% of patients have ophthalmologic phenotypes, only 2/27 (7.4%) of a cohort with complete eye examination described are described to have coloboma (Greenberg *et al.*, 1991). Paternal DNA was unavailable, however maternal DNA was negative for the c.725G>A *LLGL1* variant suggesting this variant allele was possibly paternally inherited in this patient with a maternally deleted chromosome 17p11.2. SMS is usually due to a common recurrent 3.7 Mb deletion and multiple patients with the identical deletion can show variability of expression of disease (Potocki *et al.*, 2003); however, few patients have sensorineural deafness. One patient with a more severe sensorineural deafness phenotype was found to have a point mutation in the recessive deafness gene, *MYO15A*, mapping to the critical SMS interval (Liburd *et al.*, 2001). BAB701's head circumference was 57cm at 16 years of age, which is not consistent with microcephaly. Previous head circumference measurements could not be obtained.

**BAB4807 and BAB4810**

BAB4807 was noted to be a high-risk fetus after prenatal screening revealed dilation of the lateral ventricles to 9.5mm. At 18 months, he had no speech and was crawling signifying developmental delay. He had a phenotype that included dolichocephaly, frontal bossing, simple posteriorly rotated ears, self-mutilation, micrognathia with irregular teeth, and strabismus. Cranial MRI revealed an Arnold-Chiari Type 1 Malformation. BAB4810 similarly had developmental delay, crawling at 18 months and walking at 3 years old. At 4 years of age, she was able to use two syllable words. Her phenotype included intellectual disability, frontal bossing, simple posteriorly rotated ears, and abnormal dentition. A homozygous variant in

*LLGL1* (c.2945C>T;p.A982V) was found in both affected probands (**Table S2**). An unaffected sibling and both parents were heterozygous for the variant (**Figure S7**).

**BGL – 1**

Referral diagnosis: Microcephaly, intrauterine growth restriction, radial microbrain and immature gyral pattern.

**BGL – 2**

Referral diagnosis: Global developmental delay with aphasia, microcephaly and seizures, progressive weakness, lethargy with episodes of irregular breathing and respiratory acidosis, MRI showing progressive diffuse central white matter atrophy and gliosis, hypomyelination, atrophy of thalami and pons. (**Table S2**, **Figure S1**)

**BGL – 3**

Referral diagnosis: Intellectual disability, ataxia, spasticity, autism spectrum disorder, speech delay, and anxiety disorder. (**Table S2, Figure S1**)

**BGL – 4**

Referral diagnosis: Spinal muscular atrophy complicated by failure to thrive and respiratory insufficiency, microcephaly, short stature, scoliosis, delayed motor milestones, progressive weakness, hypotonia, history of prematurity and intrauterine growth retardation (**Table S2, Figure S5).**

**BGL – 5**

Referral diagnosis: Prematurity, spasticity of inferior limbs, seizures, contractures of the Achilles, myopia (**Table S2, Figure S5).**

Table S1. Summary of reported *ANKLE2* and *VRK1* cases.

| Family ID† | Age‡ | Proband ID | Variant | Zyg | Microcephaly | Brain Abnormalities | Additional Findings | Study |
|---|---|---|---|---|---|---|---|---|
| **Yamamoto -1** | Birth | LR06-300a1 | *ANKLE2 c.1717C>G;p.L573V c.2344C>T;p.Q782** | Comp. Het | -9SD | Agenesis of corpus callosum, small frontal horns and enlarged posterior horns of the lateral ventricles, simplified gyral pattern with mildly thickened cortex | Low forehead, scalp rugae, spasticity, multiple hyper/hypo-pigmented macules | Yamamoto et al. |
| **Yamamoto -1** | Birth | Sister of LR06-300a1 | *ANKLE2* c.1717C>G;p.L573V c.2344C>T;p.Q782* | Comp. Het | + (Z-score not noted) | NR | Spasticity, multiple hyper/hypo-pigmented macules | Yamamoto *et al*. |
| **88** | Birth | 13DG0559 | *ANKLE2* c.1754G>T;p.G585V | Hom | -2.6SD | Agenesis of posterior corpus callosum | No additional abnormalities noted | Shaheen *et al.* |
| **Shaheen-1** | Birth | MC22901 | *ANKLE2* c.601G>T;p.G201W | Hom | -6SD | No additional abnormalities noted | ID, hyperactivity, sleep-cycle disturbance | Shaheen *et al*. |
| **Shaheen-2** | Birth | UCSF Case | *ANKLE2* c.601G>T;p.G201W | Hom | -4.33SD | Microlissencephaly of the cerebellum, vermian hypoplasia, partial corpus callosum agenesis, colpocephaly, pachygyria | Sloping forehead, overriding sutures, micrognathia, strabismus, high/broad nasal bridge, high-arched palate, bulbous nasal tip, ASD and ADHD | Shaheen *et al*. |
| **Link-1** | NA | LR17-511 | *ANKLE2* c.686T>G;p.V229G | Hom | -8SD | + (Not specified) | Skin pigment abnormality | This study |

| | | | | | | | | |
|---|---|---|---|---|---|---|---|---|
| **Link-1** | NA | Sibling of LR17-511 | *ANKLE2* c.686T>G;p.V229G | Hom | -8SD | + (Not specified) | Skin pigment abnormality | This study |
| **Link-2** | NA | LR18-033 | *ANKLE2* c.325G>C;p.A109P c.1421-1G>C;Splicing | Comp. Het | -6SD | + (Not specified) | No additional abnormalities noted | This study |
| **Renbaum-1** | Birth | IV-12 | *VRK1* c.1072C>T;p.R358* | Hom | -6SD | Pontocerebellar hypoplasia | Motor and sensory neuropathy | Renbaum *et al*. |
| **M017N** | NA | M017N-1 | *VRK1* c.397C>T;p.R133C | Hom | NA | Pontocerebellar hypoplasia | Pontocerebellar Hypoplasia Type 1 | Najmabadi *et al*. |
| **M017N** | NA | M017N-2 | *VRK1* c.397C>T;p.R133C | Hom | NA | Pontocerebellar hypoplasia | Pontocerebellar Hypoplasia Type 1 | Najmabadi *et al*. |
| **M017N** | NA | M017N-3 | *VRK1* c.397C>T;p.R133C | Hom | NA | Pontocerebellar hypoplasia | Pontocerebellar Hypoplasia Type 1 | Najmabadi *et al*. |
| **M017N** | NA | M017N-4 | *VRK1* c.397C>T;p.R133C | Hom | NA | Pontocerebellar hypoplasia | Pontocerebellar Hypoplasia Type 1 | Najmabadi *et al*. |
| **HOU 1183** | In utero | BAB3280 | *VRK1* c.706G>A;p.V236M c.266G>A;p.R89Q | Comp. Het | ~ -4SD | Simplified gyral pattern | Motor and sensory neuropathy | Gonzaga-Jauregui *et al*. |
| **HOU 1183** | 1y | BAB3022 | *VRK1* c.706G>A;p.V236M c.266G>A;p.R89Q | Comp. Het | ~ -6SD | Simplified gyral pattern | Motor and sensory neuropathy | Gonzaga-Jauregui *et al*. |
| **HOU 2076** | In utero | BAB5311 | *VRK1* c.1072C>T;p.R358* | Hom | ~ -6SD | Simplified gyral pattern, underdeveloped vermis | Motor and sensory neuropathy | Gonzaga-Jauregui *et al*. |
| **Nguyen-1** | 27y | 9320872 | *VRK1* c.356A>G;p.H119R c.961C>T;p.R321C | Comp. Het | NA | No abnormalities noted | Amyotrophic Lateral Sclerosis | Nguyen *et al*. |
| **Stoll-1** | Late teens | II:1 | *VRK1* c.356A>G;p.H119R c.1072C>T;p.R358* | Comp. Het | NA | Mild to moderate generalized atrophy | Motor Neuropathy | Stoll *et al*. |
| **Stoll-1** | 15y | II:4 | *VRK1* c.356A>G;p.H119R c.1072C>T;p.R358* | Comp. Het | NA | Mild to moderate generalized atrophy | Motor Neuropathy | Stoll *et al*. |

| | | | | | | | | |
|---|---|---|---|---|---|---|---|---|
| **Stoll-2** | 3y | III:4 | *VRK1* c.403G>A;p.G135R c.583T>G;p.L195V | Comp. Het | + (Z-score not noted) | No abnormalities noted | Juvenile Amyotrophic Lateral Sclerosis with Sensory Neuropathy | Stoll *et al*. |
| **78** | NA | 10DG1904 | *VRK1* c.236C>T;p.P79L | Hom | -5.1SD | Delayed myelination, posterior corpus callosum agenesis, mild substance reduction in the posterior fossa | Dysarthria, global developmental delay | Shaheen *et al*. |
| **79** | NA | 11DG0443 | *VRK1* c.236C>T;p.P79L | Hom | NA | Pachygyria, Dandy Walker malformation | Prominent nasal tip, thin upper lip, large ears, global developmental delay, talipes equinovarus, scoliosis, seizures | Shaheen *et al*. |
| **Feng-1** | 15y | V-I | *VRK1* c.1124G>A;p.W375* | Hom | No microcephaly | No abnormalities noted | Motor neuropathy | Feng *et al*. |

†If no family ID was provided, first author of study along with a number was used to denote family.
‡At presentation to medical attention or when symptoms were first noted.

Table S2. Summary of ANKLE2 pathway, PAR complex, and LLGL1 encoding variants in BGL and BHCMG databases.

| Family ID† | Proband ID | Variant | Zyg | Chr:Pos | vR/tR | phyloP | ARIC | CADD | gnomAD Het/Hom | AOH block around gene |
|---|---|---|---|---|---|---|---|---|---|---|
| **BGL-1** | Case 1 | *ANKLE2* c.706C>T;p.R236* | Compound | 12:133327370 | 71/131 | 2.963 | NR | 38 | 1/0 | NA |

| | | | | | | | | | | |
|---|---|---|---|---|---|---|---|---|---|---|
| | | c.1606C>T;p.R536C | Het | 12:133312086 | 51/103 | 3.499 | NR | 26.3 | 2/0 | |
| **BGL-2** | Case 2 | *ANKLE2*<br>c.23C>T;p.A8V<br>c.80C>G;p.A27G | Compound<br>Het | 12:133338362<br>12:133338305 | 7/25<br>20/38 | -0.023<br>1.17 | NR<br>NR | 17.45<br>24.7 | 1/0<br>0/0 | NA |
| **BGL-3** | Case 3 | *ANKLE2*<br>c.1522C>T;p.R508C<br>c.1907C>T;p.S636F | Compound<br>Het | 12:133313550<br>12:133306841 | NR<br>NR | 1.209<br>1.246 | NR<br>NR | 15.26<br>26.6 | 12/0<br>2/0 | NA |
| **BGL-4** | Case 4 | *VRK1*<br>c.266G>A;p.R89Q<br>c.706G>A;p.V236M | Phasing<br>Unavailable | 14:97312481<br>14:97321690 | NR<br>NR | 5.996<br>6.364 | NR<br>NR | 25.9<br>31 | 8/0<br>6/0 | NA |
| **BGL-5** | Case 5 | *VRK1*<br>c.3G>A;p.M1_D31del<br>c.614G>A;p.C205Y | Phasing<br>Unavailable | 14:97299811<br>14:97321598 | NR<br>NR | 4.683<br>4.203 | NR<br>NR | 26.8<br>29.3 | 0/0<br>0/0 | NA |
| **HOU 1939** | BAB4821 | *VRK2*<br>c.1234G>A;p.D412N | Hom | 2:58386604 | 60/61 | 2.252 | 0 | 24.2 | 5/0 | 7.7Mb |
| **HOU 2812** | BAB7812 | *VRK3*<br>c.139+2T>G;Splicing | Hom | 19:50519279 | 44/44 | 2.252 | 0 | 22.8 | 0/0 | 4.4Mb |
| **HOU 2812** | BAB7812 | *RTN2*<br>c.938dupC;p.P313fs | Hom | 19:45996512 | 66/74 | NA | 0 | NA | 11/0 | 5.6Mb |
| **HOU 2812** | BAB7812 | *PNKP*<br>c.931C>T;p.R311C | Hom | 19:50365800 | 115/115 | 2.395 | 0 | 34 | 1/0 | 4.4Mb |
| **HOU 3871** | BAB10531 | *PARD3B*<br>c.1222G>A;p.G408S | Hom | 2:205989107 | 91/91 | 0.557 | 11 | 34 | 52/0 | 22Mb |
| **HOU 3010** | BAB8223 | *PARD3B*<br>c.1654G>A;p.A552T | Hom | 2:206036968 | 42/42 | 2.665 | 0 | 33 | 7/0 | 37Mb |
| **HOU 234** | BAB701 | *LLGL1*<br>c.725G>A;p.S242N | Hemizygous | 17:18137597 | 38/38 | 2.706 | 0 | 24.8 | 0/0 | NA |
| **HOU 1936** | BAB4807 | *LLGL1*<br>c.2945C>T;p.A982V | Hom | 17:18145542 | 91/92 | 0.63 | 0 | 18.18 | 3/0 | 2.2Mb |
| **HOU 1936** | BAB4810 | *LLGL1*<br>c.2945C>T;p.A982V | Hom | 17:18145542 | 96/97 | 0.63 | 0 | 18.18 | 3/0 | 2.2Mb |

†For Baylor Genetics Laboratories cases, BGL-# was used to denote family.

Figure S1

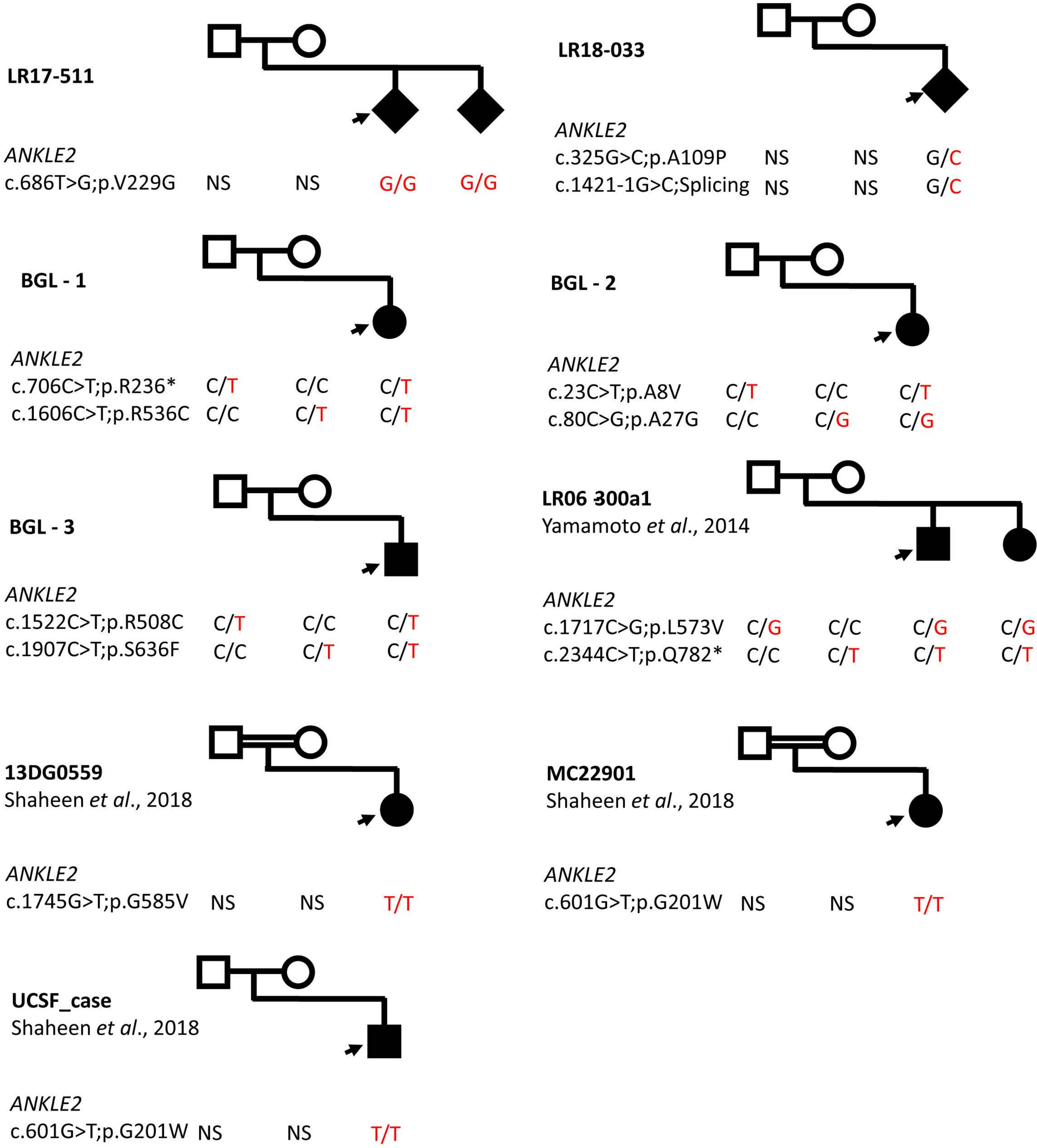
LR17-511
ANKLE2
c.686T>G;p.V229G NS NS G/G G/G
LR18-033
ANKLE2
c.325G>C;p.A109P NS NS G/C
c.1421-1G>C;Splicing NS NS G/C
BGL - 1
ANKLE2
c.706C>T;p.R236* C/T C/C C/T
c.1606C>T;p.R536C C/C C/T C/T
BGL - 2
ANKLE2
c.23C>T;p.A8V C/T C/C C/T
c.80C>G;p.A27G C/C C/G C/G
BGL - 3
ANKLE2
c.1522C>T;p.R508C C/T C/C C/T
c.1907C>T;p.S636F C/C C/T C/T
LR06-300a1
Yamamoto et al., 2014
ANKLE2
c.1717C>G;p.L573V C/G C/C C/G C/G
c.2344C>T;p.Q782* C/C C/T C/T C/T
13DG0559
Shaheen et al., 2018
ANKLE2
c.1745G>T;p.G585V NS NS T/T
MC22901
Shaheen et al., 2018
ANKLE2
c.601G>T;p.G201W NS NS T/T
UCSF_case
Shaheen et al., 2018
ANKLE2
c.601G>T;p.G201W NS NS T/T

Figure S2

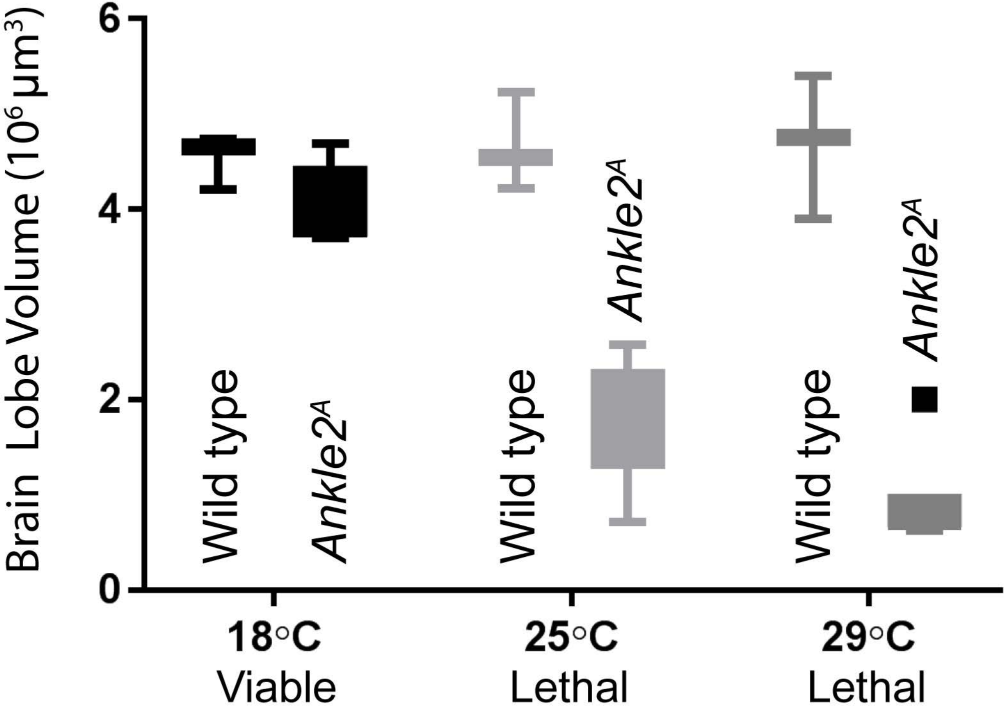

Brain Lobe Volume (10⁶ μm³)
6
4
2
0
Wild type
Ankle2A
Wild type
Ankle2A
Wild type
Ankle2A
18°C
Viable
25°C
Lethal
29°C
Lethal

Figure S3

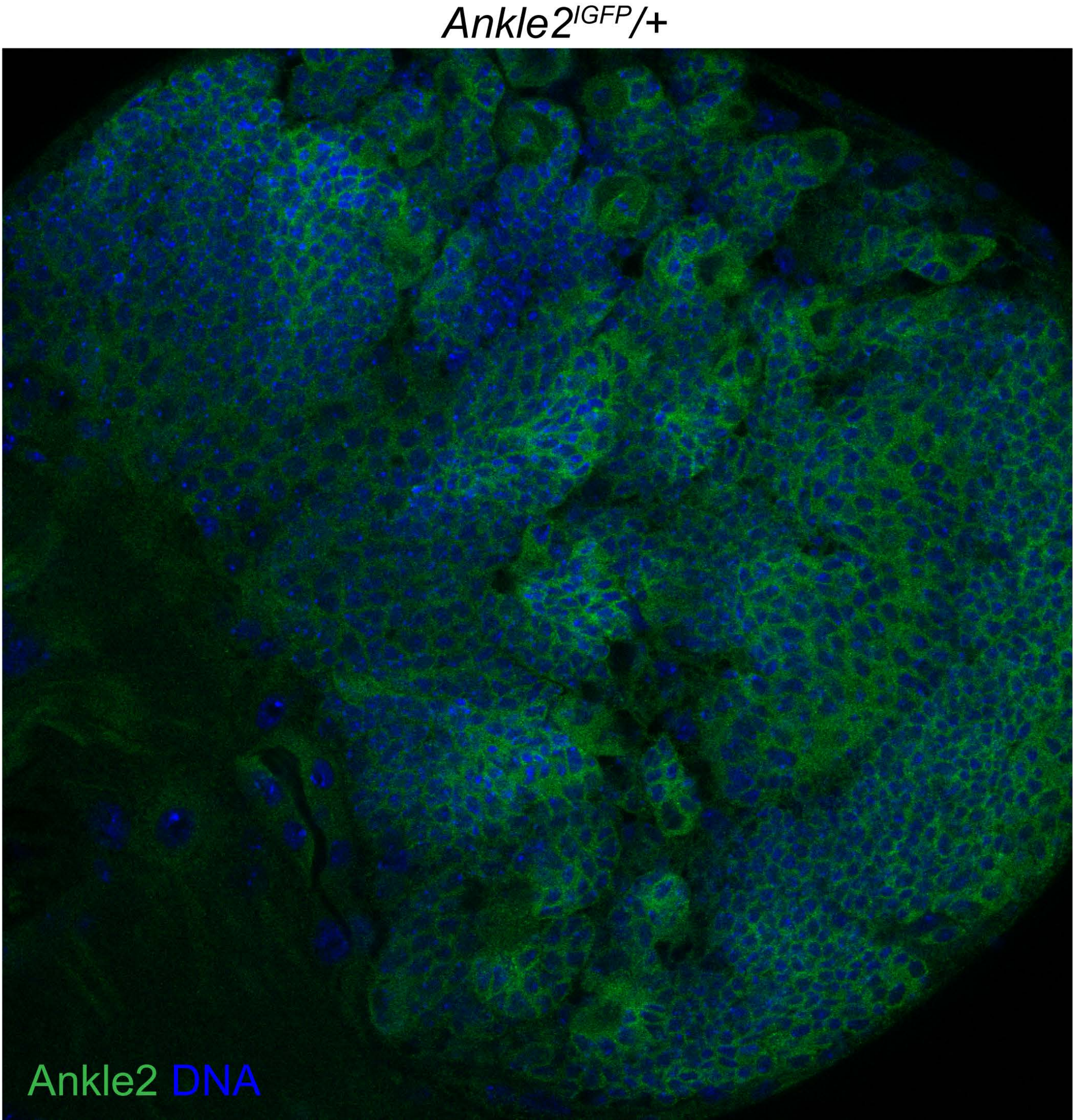

Figure S4

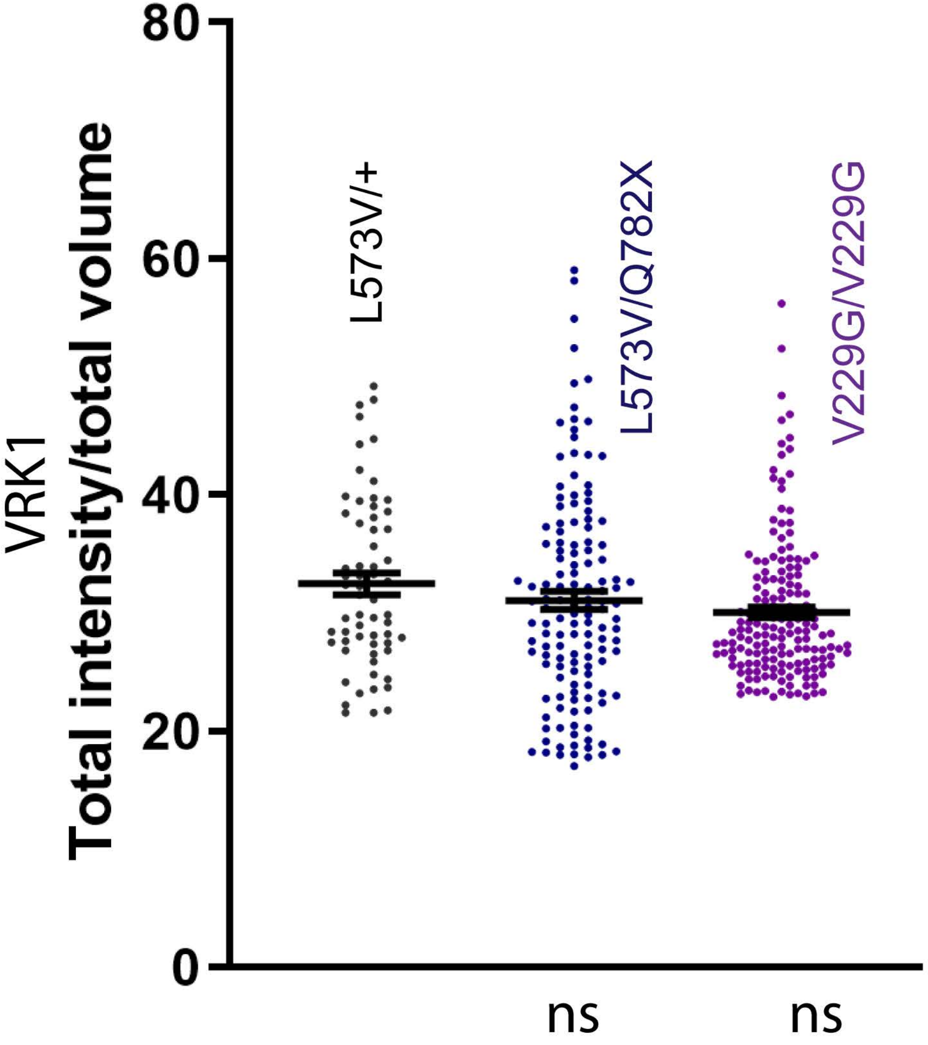

VRK1
Total intensity/total volume
80
60
40
20
0
L573V/+
L573V/Q782X
V229G/V229G
ns
ns

Figure S5

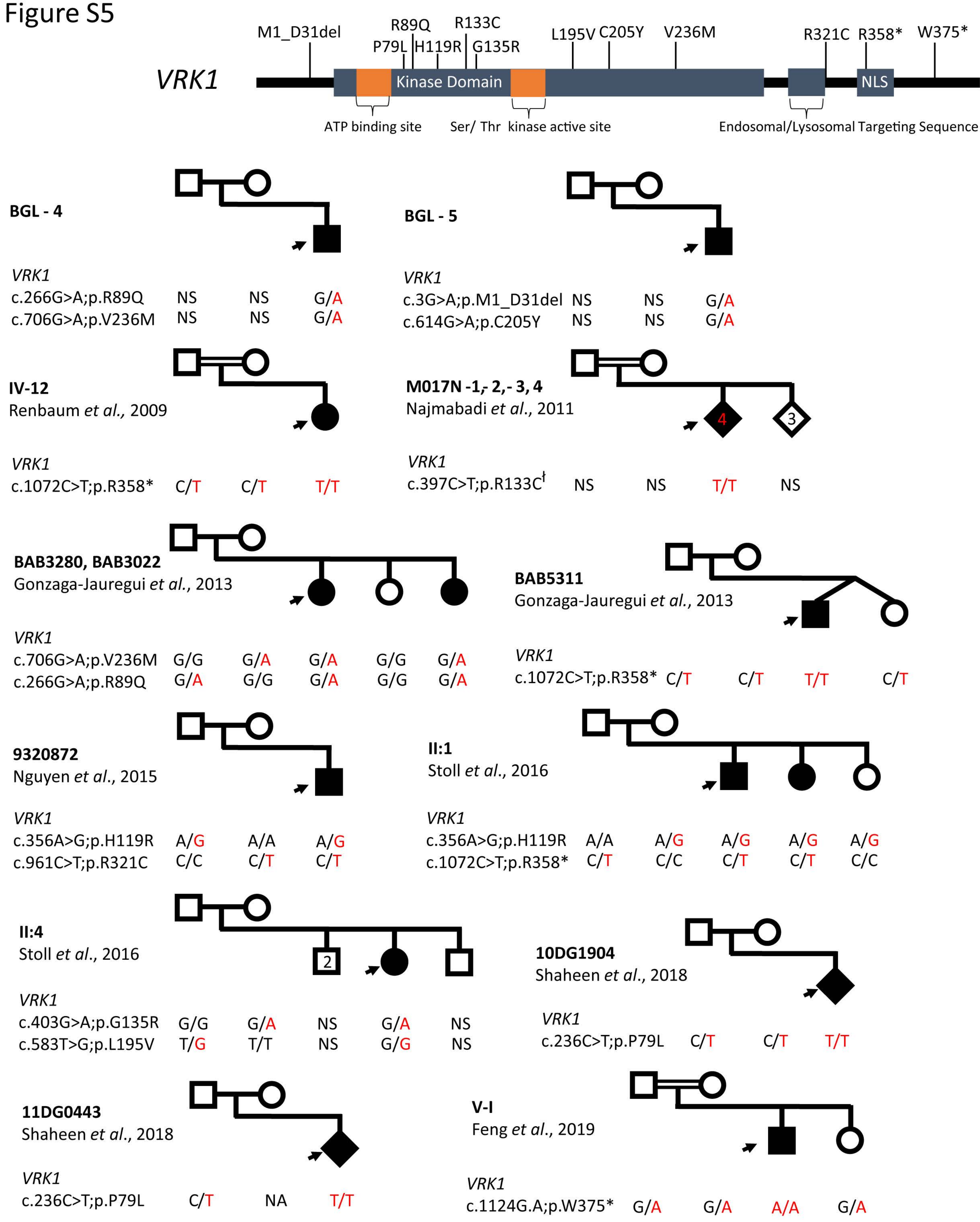


[ł]7 individuals genotyped and variants segregate with disease, however genotyped individuals other than affected individuals not specified

Figure S6

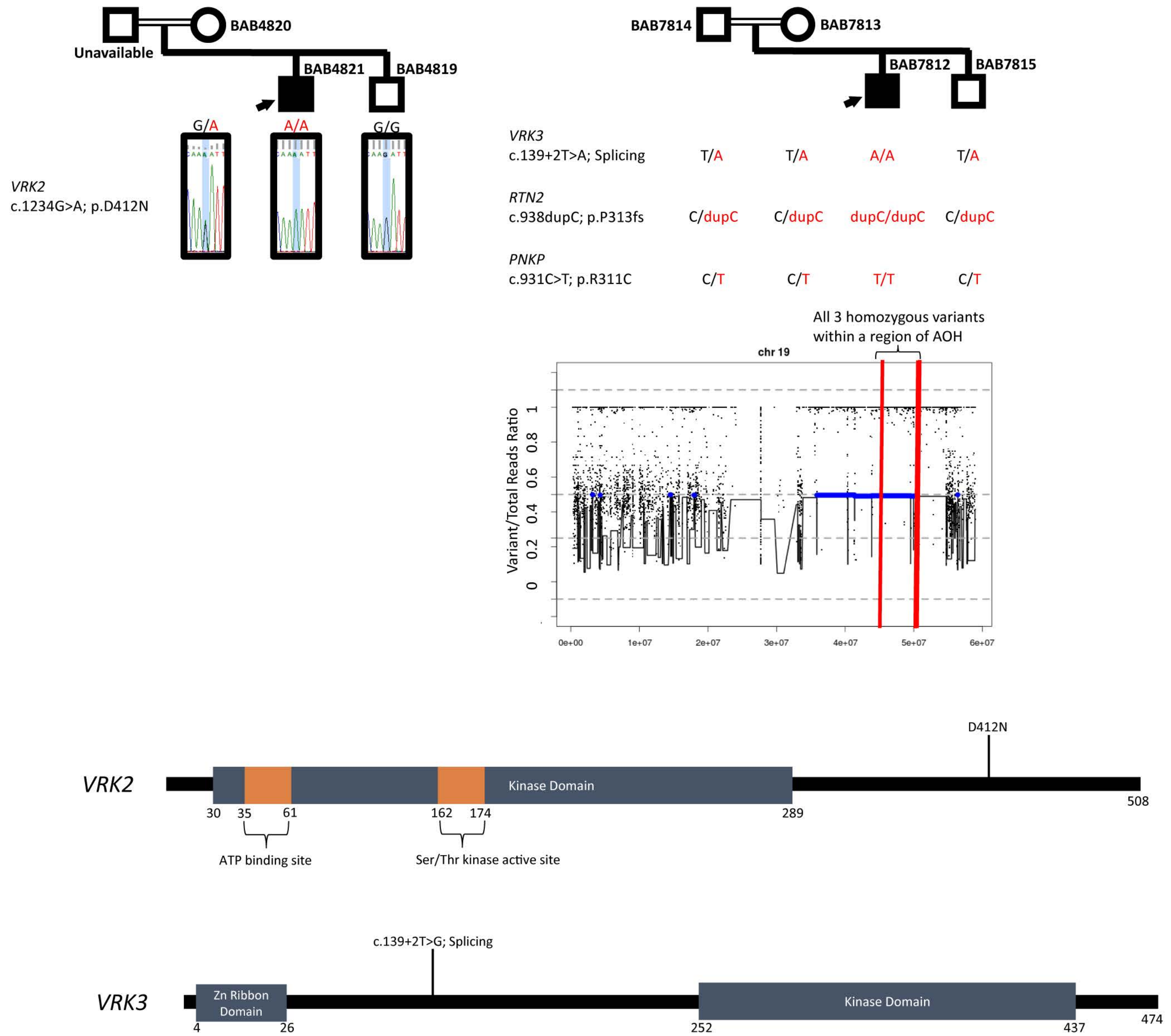
BAB4820
Unavailable
BAB4821
BAB4819
G/A
A/A
G/G
VRK2
c.1234G>A; p.D412N
BAB7814
BAB7813
BAB7812
BAB7815
VRK3
c.139+2T>A; Splicing
T/A
T/A
A/A
T/A
RTN2
c.938dupC; p.P313fs
C/dupC
C/dupC
dupC/dupC
C/dupC
PNKP
c.931C>T; p.R311C
C/T
C/T
T/T
C/T
All 3 homozygous variants within a region of AOH
chr 19
Variant/Total Reads Ratio
0
0.2
0.4
0.6
0.8
1
0e+00
1e+07
2e+07
3e+07
4e+07
5e+07
6e+07
D412N
VRK2
Kinase Domain
30
35
61
162
174
289
508
ATP binding site
Ser/Thr kinase active site
c.139+2T>G; Splicing
VRK3
Zn Ribbon Domain
Kinase Domain
4
26
252
437
474

### Figure S7

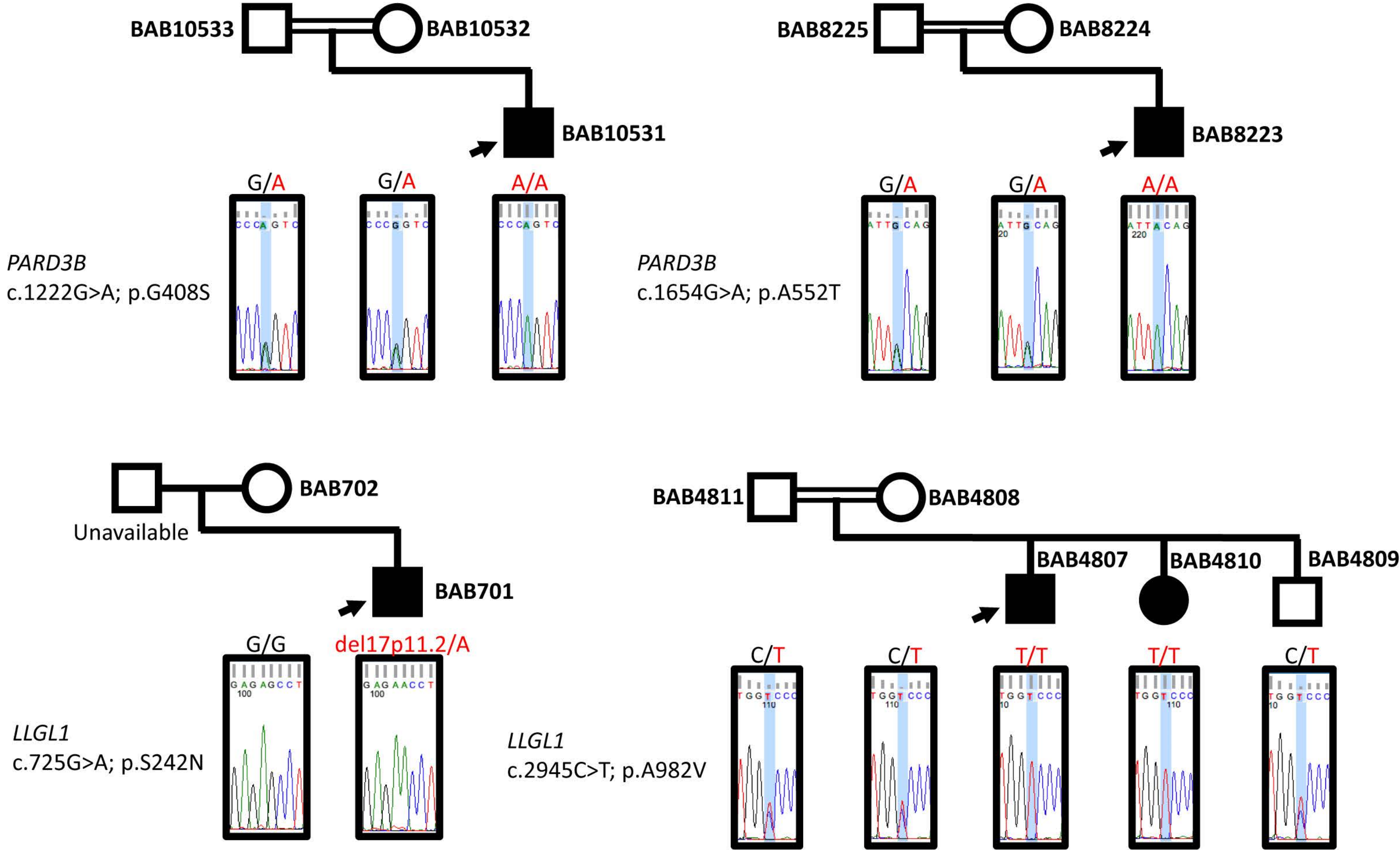

BAB10533
BAB10532
BAB10531
G/A
G/A
A/A
*PARD3B*
c.1222G>A; p.G408S
BAB8225
BAB8224
BAB8223
G/A
G/A
A/A
*PARD3B*
c.1654G>A; p.A552T
BAB702
Unavailable
BAB701
G/G
del17p11.2/A
*LLGL1*
c.725G>A; p.S242N
BAB4811
BAB4808
BAB4807
BAB4810
BAB4809
C/T
C/T
T/T
T/T
C/T
*LLGL1*
c.2945C>T; p.A982V
